# MATR3-antisense LINE1 RNA meshwork scaffolds higher-order chromatin organization

**DOI:** 10.1101/2022.09.13.506124

**Authors:** Yuwen Zhang, Xuan Cao, Zehua Gao, Xuying Ma, Qianfeng Wang, Xiumei Cai, Yan Zhang, Zhao Zhang, Gang Wei, Bo Wen

## Abstract

Long interspersed nuclear elements (LINEs) play essential role in shaping chromatin state, while the factors that cooperate with LINEs and their roles in higher-order chromatin organization remain poorly understood. Here we show that MATR3, a nuclear matrix protein, interplays with antisense LINE1 (AS L1) RNAs to form into a gel-like meshwork via phase-separation, providing a partially dynamic platform for chromatin spatial organization. Either depletion of MATR3 or AS L1 RNAs changes nuclear distribution of each other and leads to chromatin reorganization in the nucleus. After MATR3 depletion, topologically associating domains (TADs) that highly transcribed MATR3-associated AS L1 RNAs showed a decrease on local chromatin interactions. Furthermore, amyotrophic lateral sclerosis (ALS)-associated MATR3 mutants alter biophysical features of the MATR3-AS L1 RNA meshwork and cause chromatin reorganization. Collectively, we revealed an essential role of meshwork formed by nuclear matrix and retrotransposon-derived RNAs in gathering chromatin in the nucleus.

## Main

Non-coding RNAs (ncRNAs) can act as structural molecules participating in genome organization, mostly through interacting with the chromatin near their transcription loci^1^. RNA-binding proteins (RBPs) contribute to the *in-cis* interaction, and ncRNAs may promote liquid-liquid phase separation (LLPS) of RBPs, further facilitating the chromatin compaction^1–4^. Besides conventional ncRNAs, repeat element derived RNAs play important roles in the organization of higher-order chromatin architecture. For example, telomeric repeat containing RNA TERRA and major satellites (MajSAT) RNAs are essential for the higher-order organization of telomeres^5^ and the pericentric heterochromatin^6^, respectively. Furthermore, C_0_T-1 repeat RNAs (including LINEs and SINEs) associate with euchromatin and the nascent repeat-rich RNAs function as scaffolds countering DNA compaction^7, 8^.

LINE1(L1) repeat elements comprise 17% and 19% genomic region in human and mouse, respectively^9, 10^. Based on the observation that the X chromosome has higher L1 composition than autosomes, it suggested the role of L1 elements as “boosters” during X-chromosome inactivation (XI)^11–13^. Further investigations showed that silent LINEs are involved in heterochromatin compartmentalization at early XI, and the actively-transcribed LINEs (young LINEs) help to spread the inactive state to escape-prone regions, promoting facultative heterochromatin compaction at late stage of XI^14, 15^. L1 repeat RNAs were shown to interact with the chromatin domains from where they were transcribed^7, 16^. Functionally, sense-transcribed L1 RNAs can act as the nuclear scaffold, recruiting Nucleolin/KAP1 proteins to inactivate 2C-related Dux-loci^17^. Further studies showed that N6-methyladenosine (m6A) marks on these chromatin-associated L1 RNAs help regulating chromatin accessibility by recruiting m6A reader YTHDC1 or m6A eraser FTO, further affecting histone modification on nearby chromatin^18–20^. As for L1 RNAs in higher-order chromatin organization, a recent study demonstrated that L1 elements and their transcripts may instruct the segregation of inactive B compartments, and depletion of sense-transcribed L1 RNAs in ESCs disrupted the homotypic contacts between L1-rich chromosome regions^21^; CBX5/HP1α and L1 RNAs may phase-separate into larger liquid droplets and promote heterochromatin compaction^21^. Of note, L1 elements and their transcripts are widespread distributed in nucleus, while HP1α proteins mainly distribute as large foci in the nucleus, suggesting other factors may also work with L1 RNAs in chromatin structure organization.

In this study, we revealed functional roles of antisense-transcribed L1 (AS L1) RNAs and a nuclear matrix (NM) protein Matrin-3 (MATR3) in chromatin organization. NM is a network-like nuclear structure proposed as the platform for various functions in the nucleus^22, 23^. Although it has long been hypothesized that NM scaffolds the chromatin organization^24, 25^, the roles of NM components in 3D genome organization just began to be revealed. Recently, we demonstrated that NM proteins SAF-A/HNRNPU and SAFB can regulate the higher-order organization of euchromatin and heterochromatin at the genome-wide scale, respectively^6, 26^. Interestingly, the functional roles of SAF-A/HNRNPU and SAFB in chromatin organization were dependent on chromatin-associated RNAs^6, 27^. MATR3 is one of the first identified NM components that presents unique physicochemical properties^28^. Functionally, MATR3 is implicated in pre-mRNA splicing^29–31^ and affect biological processes including pluripotency maintenance^32, 33^ and X chromosome inactivation (XI) ^2, 34^. A recent 3D genome study in erythroid cells showed that MATR3 stabilizes the chromatin occupancy of CTCF and cohesin at a subset of sites; loss of MATR3 affects weak-insulated topologically associating domains (TADs) and accelerates cell fate transition^35^. Here we show that MATR3 mediates chromatin interaction by interacting with AS L1 RNAs, which may enhance our understandings of repeat RNAs in chromatin organization.

## Results

### MATR3 regulates the spatial organization of nuclear chromatin

To investigate the fine scale localization pattern of MATR3 in the nucleus, we performed super-resolution fluorescence microscopy and immuno-electron microscopy in mouse hepatocytes (AML12 cells), which had been applied in studying nuclear architecture previously^6, 26^. The imaging data showed that MATR3 proteins organize into network-like structures with some concentrated puncta, which adjacent to chromatin fibers in the nucleus (Fig. 1a and Extended Data Fig. 1a). To determine the chromatin types associated with MATR3, we measured the correlation coefficient (r) between MATR3 and histone marks on randomly selected regions of interest (ROI) in the nuclei. The nuclear distribution of MATR3 showed the highest correlation with H3K27me3, comparing to other histone marks including H3K9me3, H3K9me2, H3K27ac or H3K4me3 (Fig. 1b and Extended Data Fig. 1b).

**Fig. 1:**
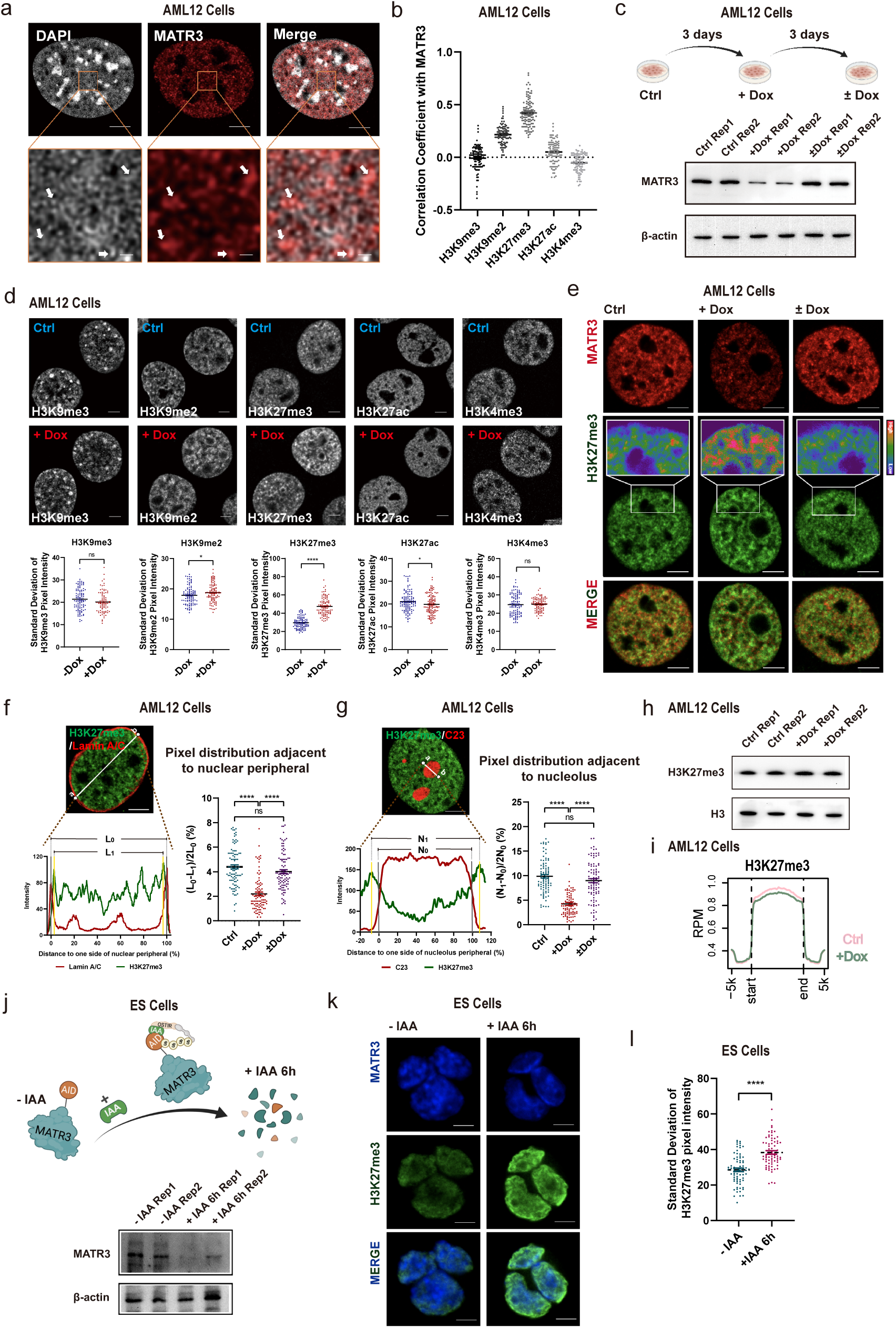
MATR3 regulates the spatial organization of chromatin. **a**, Super-resolution fluorescence microscopy images showing relative distribution between MATR3 and DAPI. **b**, Coefficient of correlation between MATR3 and histone modification H3K9me3 (n=105), H3K9me2 (n=98), H3K27me3 (n=107), H3K27ac (n=97) and H3K4me3 (n=99) in AML12 cells. Quantifications were performed on randomly selected ROIs in cell nuclei. Also see Extended Data Fig.1b. Each point represents one cell. **c**, (upper) Schematic diagram of dox-inducible shRNA system for MATR3 knockdown and MATR3 rescue in AML12 cells. (lower) Western blotting detected the expression level of MATR3 after 3 days of Dox treatment (+Dox) and followed by 3 days of Dox removal (±Dox) in AML12 cells. Rep, replicate. **d**, (upper) Representative cross-section images showing distribution of histone modifications upon Ctrl and MATR3 knock down (+Dox). (lower) Quantify the distribution pattern of histone modifications by Standard Deviation of Pixel Intensity in cell nuclei. For H3K9me3, n=102 (Ctrl) or 84 (+Dox); for H3K9me2, n=100 (Ctrl) or 101 (+Dox); for H3K27me3, n=98 (Ctrl) or 98 (+Dox); for H3K27ac, n=117 (Ctrl) or 124 (+Dox); for H3K27me3, n=98 (Ctrl) or 98 (+Dox); for H3K4me3, n=107 (Ctrl) or 97 (+Dox). Each point represents one cell. **e**, Representative cross-section images showing nuclear localization of MATR3 and H3K27me3 upon Ctrl, MATR3 knock down (+Dox) and MATR3 rescue (±Dox). **f**, Relative distribution of H3K27me3 and Lamin A/C. L_0_, region between nuclear membrane (position of nuclear membrane was determined by the X-axis of the Lamin A/C pixel peaks on both sides). L_1_, region between two H3K27me3 pixel peaks that closest to the nuclear membrane. Quantify changes of H3K27me3 distribution adjacent to nuclear peripheral in Ctrl (n=95), MATR3 knockdown (+Dox) (n=92) and MATR3 rescue (±Dox) (n=98) cells by formula of (L_0_-L_1_)/2L_0_ (%). **g**, Relative distribution of H3K27me3 and C23. N_0_, region between nucleolus membrane (position of nucleolus membrane was determined by the X-axis of the half-peaks on both sides). N_1_, region between two H3K27me3 pixel peaks that are closest to the nucleolus membrane. Quantify changes of H3K27me3 distribution adjacent to nucleolus in Ctrl (n=91), MATR3 knock down (n=91) and MATR3 rescue (n=92) cells by formula of (N_1_-N_0_)/2N_0_ (%). **h**, H3K27me3 modification level upon Ctrl and MATR3 knockdown (+Dox) as detected by western blotting. Rep, replication. **i**, Average enrichment of H3K27me3 in Ctrl and shMatr3 at peaks regions (ChIP-seq). **j**, (upper) Schematic diagram of IAA-inducible rapid protein degradation system for MATR3 in ES cells. (lower) Western blotting detection of the efficiency of MATR3 knockdown in ES cells after 6h addiction of 500μM IAA or equal-volume of alcohol (-IAA). Rep, replication. **k**, Representative cross-section images showing nuclear localization of H3K27me3 in ES cells after 6h addiction of 500μM IAA (+IAA 6h) or equal-volume of alcohol (-IAA). **l**, Standard deviation of H3K27me3 pixel intensity after 6h addiction of 500μM IAA (+IAA 6h) (n=74) or equal-volume of alcohol (-IAA) (n=74). The P values were calculated using unpaired two-tailed Student’s t test; ns, not significant, *p<0.05, ****p<0.0001. Error bars indicate mean ± s.e.m. Scale bars, 5μm (**a** (upper), **e-g** and **k**) or 0.5μm (**a** (lower)).

To reveal the biological function of MATR3 proteins in chromatin regulation, we established a doxycycline (Dox)-inducible, short hairpin RNA (shRNA)-based RNAi system^36^ in AML12 cells. Using this system, three days of Dox treatment can generate a knockdown of MATR3, and three days after Dox removal can restore the expression level consistent to the control group (Fig. 1c). We first investigated the chromatin changes within regions of different chromatin types by immunofluorescence staining. To quantify their distribution pattern, we measured the standard deviation (SD) values of pixel intensity in the whole nuclear region. After MATR3 depletion, SD values of H3K9me2 and H3K27me3 increased, and that of H3K27ac decreased. Among these histone marks, H3K27me3 showed the most significant redistribution (p<0.0001) (Fig. 1d). For the following investigation, therefore, we took H3K27me3 as the representative mark to probe chromatin organization changes. H3K27me3 staining in the control group showed a relatively diffused distribution pattern with some irregular foci in the nucleus. After depletion of MATR3, H3K27me3 staining presented larger and brighter foci in the inner nucleus as well as near the nuclear periphery and the nucleolus (Fig. 1e and Extended Data Fig. 1c). To quantify the redistribution of H3K27me3-modified chromatin towards nuclear periphery or nucleolus, we analyzed the pixel signal distribution on randomly selected ROIs in Lamin A/C or C23 (a nucleolus marker) co-stained cells. The data indicated that H3K27me3-modified chromatin became significantly closer to the nuclear periphery and the nucleolus after MATR3 depletion (Fig. 1f-g and Extended Data Fig. 1f-g). Importantly, Dox removal for three days restored the H3K27me3 pixel intensity SD values and spatial distribution relative to nuclear periphery and nucleolus (Fig. 1e-g and Extended Data Fig. 1d,f,g), suggesting a direct regulatory role of MATR3 on chromatin organization. Next, we asked whether the changes of H3K27me3 staining were due to the alteration of H3K27me3 modification levels. However, the total H3K27me3 level and the genome-wide profiles were largely unchanged upon MTAR3 depletion, as indicated by western blotting (Fig. 1h) and ChIP-seq (Fig. 1i and Extended Data Fig. 1e) in AML12 cells. Therefore, the changes of H3K27me3 staining reflected the spatial redistribution of chromatin rather than the alteration of histone modification levels.

To rule out the potential off-target of RNAi methods and other secondary effects after knockdown, we established an auxin (IAA)-inducible rapid protein degradation system^37^ in mouse embryonic stem cells (ESCs), which can generate MATR3 degradation within 6 hours (Fig. 1j, Extended Data Fig. 1j). As AML12 cells cannot be cultured in clones, the acute MATR3 degradation is technically difficult in this cell line. In ES cells, we first evaluated co-localization between MATR3 and different types of chromatins. H3K27me3-modified chromatin showed positive correlation with MATR3, although the correlation in ES cells was weaker than that in AML12 cells (Extended Data Fig. 1h,i). After rapid degradation of MATR3, H3K27me3 in ES Cells showed larger and brighter staining near the nuclear periphery (Fig. 1k), and the SD of H3K27me3 pixel intensity significantly increased accordingly (Fig. 1l). Collectively, the acute degradation of MATR3 in ES Cells and RNAi-based MATR3 knockdown in AML12 cells resulted in similar spatial changes of chromatin as probed by H3K27me3 staining

### MATR3 interacts with nuclear RNAs including antisense LINE1 RNAs

Nuclear RNAs were shown to participate in chromatin spatial organization^38, 39^. As MATR3 acts as an RNA-binding protein^30, 31, 40^, we wondered whether MATR3’s functional role in chromatin organization is RNA-dependent. According to cell fractionation assay ^41^ followed by western blotting detection, most MATR3 proteins present in chromatin-associated fractions in control cells, but greatly delocalized from chromatin after RNA pol II inhibitor dichlororibofuranosylben-zimidazole (DRB) treatment or RNase A treatment (Fig. 2a). Interestingly, immunofluorescent images showed that treating cells with RNA transcription inhibitor (DRB or α-amanitin) or RNase A disrupted network of MATR3 proteins and led them forming into spheroidal puncta (Fig. 2b and Extended Data Fig. 2a,c). Moreover, deleting RNA recognition motifs (RRM1/2) of MATR3 proteins made MATR3 distribute as spheroidal foci, and the impact of RRM2 deletion was more prominent (Extended Data Fig. 2b). Pixel signal of MATR3 and H3K27me3 is positively correlated (r=0.66) in the control cell, but the correlation turned negative in the DRB (r=−0.36) or RNase A (r=−0.13) treated cells (Fig. 2b and Extended Data Fig. 2c). These data suggested that MATR3-chromatin association is RNA-dependent.

**Fig. 2:**
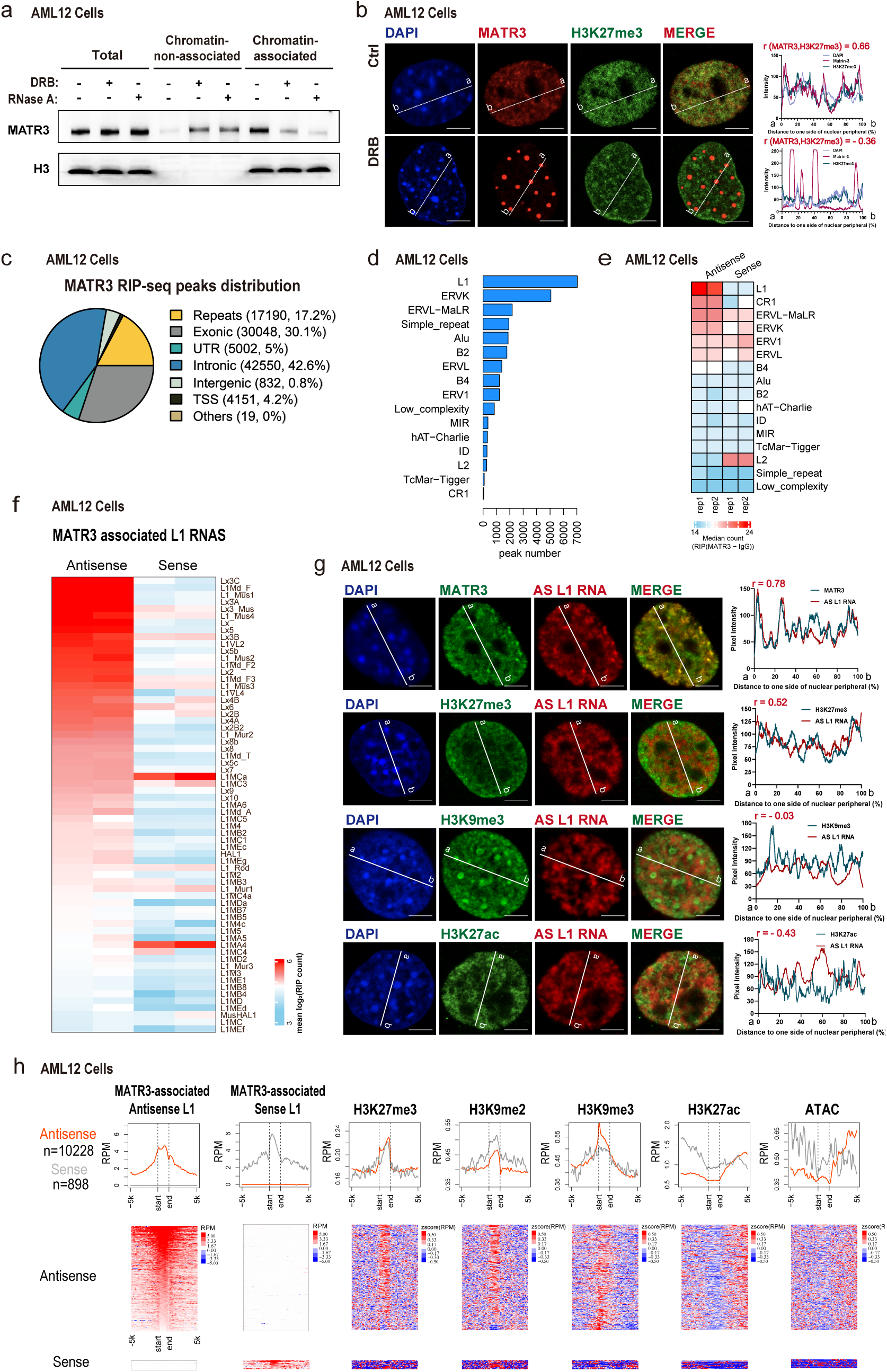
MATR3 associates with nuclear RNAs including repeat elements-derived transcripts. **a**, Western blotting showing the distribution of MATR3 proteins in chromatin-non-associated and chromatin-associated extracts before and after DRB (75μM for 12h) or RNase A (pre-treat with 0.05% Triton x-100 for 30s, followed by 10μg/ml RNase A for 1h) treatment in AML12 cells. Representative of two independent replicates with similar results. **b**, (Left) The representative cross-section image showing nuclear distribution of DAPI, MATR3 and H3K27me3 before and after 24h treating of 75μM DRB in AML12 cells. (Right) Line charts showing pixel intensity of each channel on the ROIs. r, coefficient of correlation. **c**, Genomic distribution of all MATR3 RIP-seq peaks in AML12 cells. **d**, The number of MATR3 RIP-seq peaks in repetitive elements (REs) in AML12 cells. **e**, Heatmap of MATR3 RIP-seq sense and antisense median reads count in repetitive elements in AML12 cells. All RE copies with the RIP (MATR3-IgG) count number >= 10 are kept. Median reads counts are measured for all copies of that RE family. **f**, Heatmap of RIP-seq antisense and sense mean reads count for MATR3 associated L1 subfamilies. L1 subfamilies are considered as MATR3 associated if the subfamily contains more than 50 copies. The copies with RIP (MATR3-IgG) count number >= 10 are kept. L1 RNAs are ranked by antisense mean reads count. **g**, (Left) Representative cross-section images showing relative distribution between AS L1 RNA with MATR3 and with histone modification marks (H3K27me3, H3K9me3 and H3K27ac) in AML12 cells. Probes for RNA FISH were designed towards the consensus sequence of antisense L1_Mus1 RNAs. (Right) Line charts showing pixel intensity of each channel on the ROIs. **h**, Normalized average density of the marks (top) and heatmaps(bottom) for the two groups of L1 loci that AS L1 RNA or L1 RNA interacted with MATR3. L1 loci with RIP (MATR3-IgG) count number >= 10 of antisense RNA were identified as MATR3-associated antisense L1, and the same cutoff for MATR3-associated sense L1. r, coefficient of correlation. Scale bars, 5μm (**b**, **g**).

To further identify MATR3-associated RNAs in chromatin organization, we performed strand-specific RNA immunoprecipitation sequencing (RIP-seq) in AML12 and ES cells with an anti-MATR3 antibody. In total, 42.6% RIP-seq peaks were associated with intronic regions of annotated genes in AML12 cells (Fig. 2c), which is in accordance with MATR3 PAR-CLIP data from human neuronal cells ^30^ and eCLIP data from HepG2 cells^42^. And we noticed that 17.2% of MATR3 RIP-seq peaks were distributed at repeat sequences. By quantifying each category of repetitive elements (REs) according to peak numbers and median counts, antisense (AS) L1 transcripts showed a strong association with MATR3 (Fig. 2d,e). Furthermore, for most of L1 subfamilies, the interactions between MATR3 and AS L1 RNAs were stronger than those between MATR3 and sense L1 RNAs (Fig. 2f). Our data in ES cells (Extended Data Fig. 2d) and previous MATR3 eCLIP conducted in HepG2 cells^42^ indicated the same results.

In order to visualize nuclear distribution of AS L1 RNAs, we performed fluorescent in-situ hybridization (FISH) in AML12 cells. Probes for RNA FISH were designed according to the consensus sequence of antisense transcripts from L1_Mus 1 subfamily, which ranked high on MATR3-associated AS L1 subfamilies (Fig. 2f). Remarkably, nuclear localization of AS L1 RNAs was highly correlated with that of MATR3 in both AML12 (r=0.76) (Fig. 2g) and ES (r=0.52) cells (Extended Data Fig. 2e), displaying a meshwork-like structure in the nucleus. Imaging data in AML12 cells showed that AS L1 RNAs were positively correlated with H3K27me3-modified chromatin (r=0.52), but negatively correlated with H3K27ac-modified chromatin (r=−0.43) (Fig. 2g). We then investigated the genomic features on MATR3-associated L1 elements in AML12 cells. MATR3-associated sense and AS L1 RNAs tended to be transcribed from different L1 elements, which are enriched for H3K27me3, H3K9me2 and H3K9me3 histone modifications (Fig. 2h).

Together, our genomic and imaging data revealed an extensive interaction between MATR3 and AS L1 RNAs.

### MATR3-AS L1 meshwork shapes the nuclear chromatin architecture

Next, we investigated the functional roles of the MATR3-AS L1 meshwork in higher-order chromatin organization. We tested whether AS L1 RNAs affect MATR3’s function in chromatin organization by treating AML12 cells with antisense oligonucleotides (ASOs) to knockdown AS L1 RNAs (Extended Data Fig. 2f). After 6 hours of AS L1 ASOs treatment, 22% of cells showed MATR3 forming into spheroidal foci in the nucleus; after 12 and 24 hours, this kind of cells increased to 53% and 65%, respectively. Moreover, a small portion of cells appeared spheroidal foci in both nucleus and cytoplasm (Extended Data Fig. 2g, h). Immunofluorescent staining revealed that after AS L1 knockdown, MATR3 foci disassociated from H3K27me3-modified chromatin (r=0.66 in Ctrl ASO; r=−0.11 in AS L1 ASO), resulted in a redistribution of H3K27me3 which shown as larger foci in the nucleus (Fig. 3a). Furthermore, SD of H3K27me3 pixel intensity significantly increased in AS L1 ASOs treated cells (Fig. 3b). The spatial alteration of H3K27me3 upon AS L1 depletion is similar as the phenomenon caused by MATR3 knockdown, which suggested that AS L1 RNAs may function as ‘RNA Glue’ for MATR3 proteins in maintaining the meshwork structure and their association with nuclear chromatin.

**Fig. 3:**
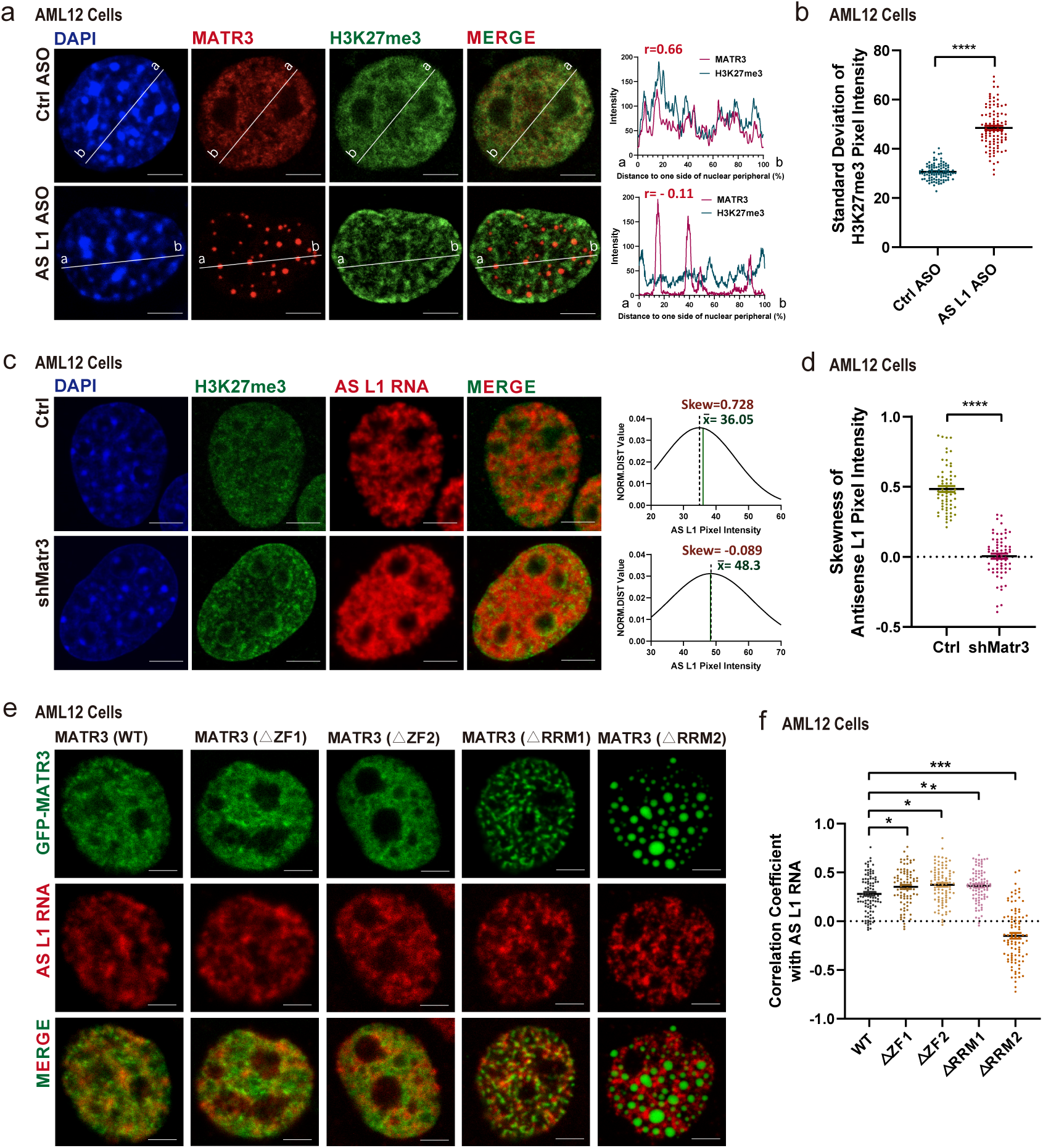
MATR3 cooperates with AS L1 RNA in chromatin organization. **a**, (Left) The representative cross-section image showing nuclear distribution of MATR3 and H3K27me3 before and after 12h treating with antisense L1 ASOs in AML12 cells. (Right) Line charts showing pixel intensity of each channel on the ROIs. r, coefficient of correlation. **b**, Standard deviation of H3K27me3 pixel intensity before (n=98) and after (n=97) 12h treating with antisense L1 ASOs in AML12 cells. **c**, (left) The representative cross-section image showing nuclear distribution of H3K27me3 and AS L1 RNA before and after MATR3 knockdown (Dox treatment for 3d) in AML12 cells. (right) The normal distribution curve for the AS L1 pixel intensity. **d**, Skewness of antisense L1 RNA pixel intensity in Ctrl (n=63) and shMatr3 cells (n=66). **e**, Representative images showing nuclear colocalization of AS L1 RNAs with wild-type and truncated GFP-MATR3 proteins in AML12 cells. **f**, Coefficient of correlation between AS L1 RNA with wild-type and truncated GFP-MATR3 proteins. WT (n=98), △ZF1 (n=83), △ZF2 (n=89), △RRM1 (n=91), △RRM2 (n=94). The P values were calculated using unpaired two-tailed Student’s t test; ns, not significant, *p<0.05, ****p<0.0001. Error bars indicate mean ± s.e.m. Scale bars, 5μm (**a**, **c**, **e**).

We further investigated whether MATR3 affects the localization of AS L1 RNAs. Immunofluorescent images showed a redistribution of AS L1 RNAs after MATR3 depletion: compared to the meshwork-like organization in control cells, AS L1 RNAs appeared a more dispersed distribution in MATR3 depleted cells (Fig. 3c). To quantify these changes, we analyzed the Gaussian fit distribution curve of the fluorescence signal. Distribution of AS L1 RNAs in control cells is right-skewed with a mean value greater than mode value and a skewness greater than zero, which suggested the presence of concentrated distribution in partial areas. After MATR3 depletion, however, the fluorescence signals of AS L1 RNAs nearly followed a standard Gaussian distribution with a mean value nearly equal to mode value and a skewness close-to-zero, which suggested a random distribution (Fig. 3c,d).

MATR3 protein has two zinc finger (ZF) domains and two RNA recognition motifs (RRMs) ^43^. We wondered which domains are necessary for MATR3 to interact with AS L1 RNAs. We expressed GFP-tagged MATR3 truncations (△ZF1, △ZF2, △RRM1 and △RRM2) in endogenous MATR3-depleted AML12 cells and examined their colocalization with AS L1 RNAs. The deletion of ZF1 or ZF2 did not affect the MATR3 distribution and its association with AS L1 RNAs. The deletion of RRM1 and RRM2 both affect MATR3 distribution, but only RRM2 deletion abolished MATR3’s association with AS L1 RNAs (Fig. 3e,f). Therefore, RRM2 domain is essential for MATR3 to bind with AS L1 RNAs.

Collectively, these results indicated that AS L1 RNAs and MATR3 affect cellular localization of each other and loss of either of them could lead to the redistribution of chromatin. AS L1 RNAs and MATR3 act together to form a meshwork-like structure that help to shape the chromatin architecture in cell nuclei.

### Phase-separation facilitates the formation of MATR3-AS L1 meshwork

To investigate the physical characters of the meshwork formed by MATR3 and AS L1 RNAs, we first tested its dynamic feature in GFP-MATR3-expressed AML12 cells with fluorescence recovery after photobleaching (FRAP) assay. After a period of 20 seconds, the bleached area recovered fluorescence intensity to half of the initial (Fig. 4a,b), indicating that the MATR3 meshwork is partially dynamic.

**Fig. 4:**
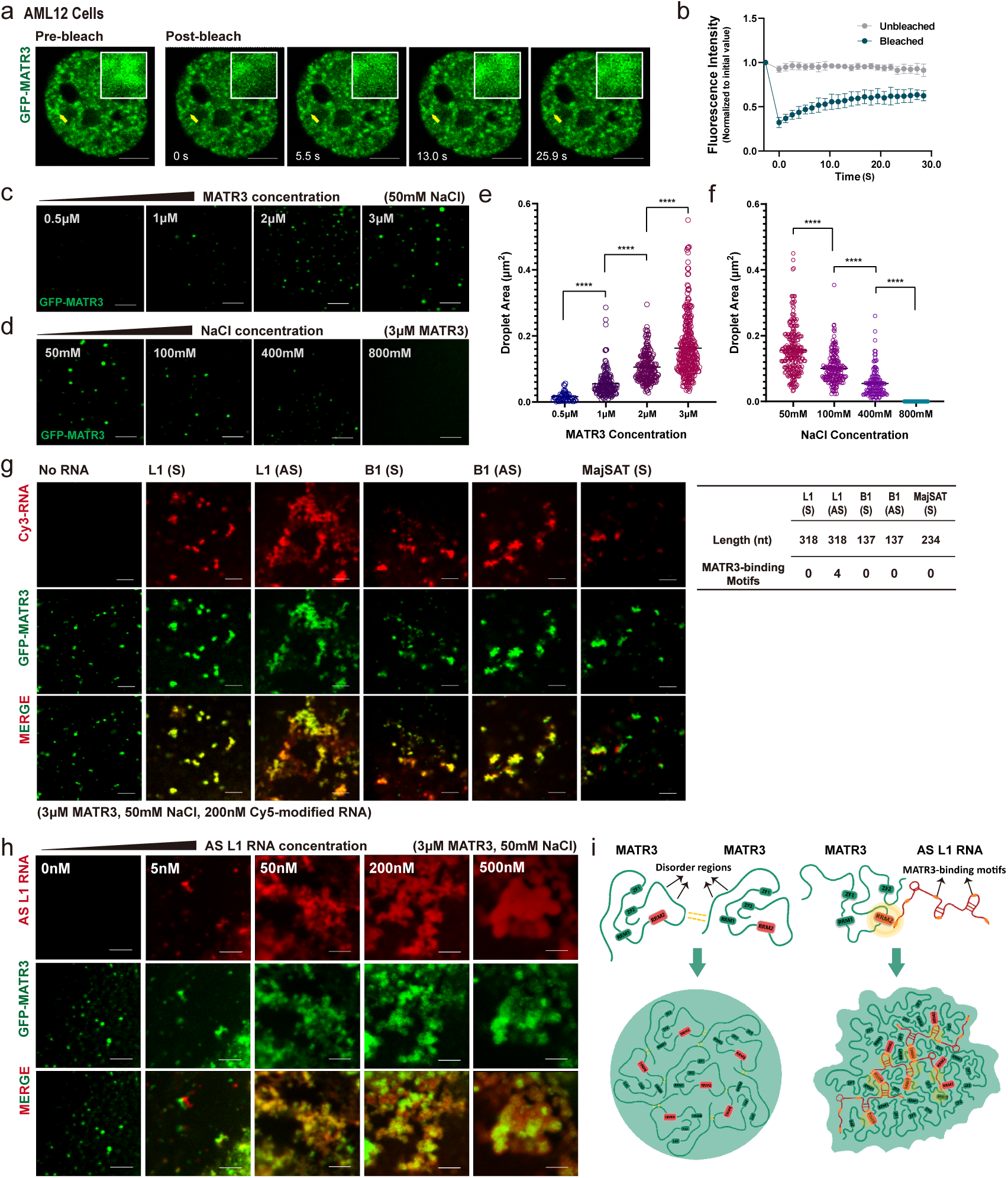
AS L1 RNAs facilitate meshwork-like organization of MATR3 proteins. **a**, Representative images of the GFP-MATR3 FRAP experiments in AML2 cells. Solid arrows indicate the bleached points. **b**, The fluorescence recovery curve of the GFP-MATR3 FRAP experiments. Data are expressed as the mean ± s.e.m. (n = 9). **c**, Representative images of droplet formation assays with different concentrations of GFP-MATR3 proteins. NaCl concentration, 50mM. **d**, Representative images of droplet formation assays with different NaCl concentrations. GFP-MATR3 protein concentration, 3 μM. **e**, Areas of MATR3 protein droplets formed in different protein concentration (3 μM: n = 315; 2 μM: n = 196; 1 μM: n = 165; 0.5 μM: n = 50). **f**, Areas of MATR3 protein droplets formed in different NaCl concentration (50 mM: n = 248; 100 mM: n = 196; 400 mM: n = 127; 800 mM: none). **g**, (left) Representative images of droplet formation assays by GFP-MATR3 with different *in-vitro*-transcribed RNAs (Sense L1, Antisense L1, Sense B1, Antisense B1). RNA concentration, 200nM. GFP-MATR3 protein concentration, 3 μM. NaCl concentration, 50mM. (right) The table showing the nucleotide number and the number of MATR3-binding motifs on the *in-vitro*-transcribed RNAs. **h**, Representative images of droplet formation assays by GFP-MATR3 with different concentration (0nM, 5nM, 50nM, 200nM, 500nM) of AS L1 RNAs. GFP-MATR3 protein concentration, 3 μM. NaCl concentration, 50mM. **i**, Schematic representation for MATR3-MATR3 droplets formation and MATR3-AS L1 RNA meshwork formation. The P values were calculated using unpaired two-tailed Student’s t test; ****p<0.0001. Scale bars, 5μm (**a**, **c**, **d**, **g**, **h**).

Recent studies have revealed dynamic nuclear compartments driven by the liquid-liquid phase separation (LLPS) of proteins and/or RNAs^44, 45^. To test the phase-separation potential of MATR3 proteins, we expressed and purified GFP-tagged MATR3 proteins and performed the *in vitro* droplet formation assay. As expected, GFP-MATR3 proteins formed spherical assemblies at room temperature; area of these assemblies increased with higher protein concentration and decreased with higher NaCl concentration (Fig. 4c-f). When incubating 3 μM of GFP-MATR3 proteins with total RNAs extracted from AML12 cells, 5-10 ng/ul of total RNAs facilitated GFP-MATR3 proteins to form larger irregular particles, while more than 20 ng/ul of total RNAs buffered these particles (Extended Data Fig. 3a). These results are in agreement with reported features of phase-separated proteins^46^.

To examine roles of repeat RNAs in shaping MATR3 condensates *in vitro*, we incubated GFP-MATR3 proteins with various *in-vitro* transcribed, Cy5-modified repeat RNAs and observed their droplet formation behaviors. The B1 and major satellites (MajSAT) RNAs were transcribed from full-length B1 or MajSAT elements, respectively. As the full-length L1 element is too long (∼6kb), the L1 RNAs were transcribed from a 318bp-consensus sequence of L1 Md_F2 elements (Extended Data Fig. 3b). Intriguingly, AS L1 RNAs facilitated MATR3 proteins forming into mesh-like assemblies *in vitro*. While under the same molecule concentration, other repeat RNAs only made MATR3 droplets to undergo a slight deformation (Fig. 4g).

To further interpret the action mechanisms of each repeat RNAs, we analyzed the numbers of MATR3-binding motifs^30^ on them. As expected, only the AS L1 sequence contains MATR3-binding motifs. Furthermore, motif density on the 318nt-antisense L1 Md_F2 RNA fragment (4/318) is comparable to that on full-length antisense L1 Md_F2 RNA (89/5948) and is much higher than that on full-length sense L1 Md_F2 RNA (4/5948) (Extended Data Fig. 3b). Hence, we suggest that AS L1 RNA has a higher affinity to bind MATR3 proteins. In cell nuclei, MATR3 could stochastically interact with different RNAs due to electrostatic forces. Those non-specifically interacting RNAs would lower MATR3’s saturation concentration and buffer the MATR3 liquids at high RNA concentration^46, 47^. While the specifically interacting RNAs like AS L1 RNAs could form multivalent interaction with MATR3 proteins, that may enhance the overall avidity^48^ and promote the meshwork-like assembly formation.

Furthermore, when incubating 3μM of MATR3 proteins with different concentrations of AS L1 RNAs, 5nM to 200nM AS L1 RNAs could help the formation of the meshwork-like structure (Fig. 4h). According to a quantitative proteomics analysis, there are in average 3,240,673 MATR3 protein molecules in each mammalian cell^49^. So, the estimated nuclear concentration of MATR3 protein is about 5.6 μM (close to 3μM MATR3 concentration within *in vitro* system). In addition, it was estimated that about 1,000 L1 RNA copies were present in one mammalian cell^39^, with a concentration around 1 nM. Considering that some cellular AS L1 RNAs are full-length transcribed (containing ∼20 copies of the 318nt-AS L1 RNA fragment) or have diverse motif density, we suggest the *in vitro* system containing 5nM to 50nM 318nt-AS L1 RNA fragment (Fig. 4h) could nearly mimic the stoichiometric ratio between MATR3 and AS L1 RNAs *in vivo*.

In summary, we suggest that MATR3 proteins have the potential to undergo liquid-liquid phase separation, the weak interaction between proteins’ disorder regions contribute to this behavior. When MATR3 proteins interact with AS L1 RNAs via the RRM2 domain, the affinity between molecules increased, thereby facilitating the higher-order MATR3-AS L1 meshwork formation (Fig. 4i).

### 3D genome organization changes upon MATR3 depletion

To study the role of MATR3 on 3D genome organization, we performed Hi-C in control and *Matr3*-depleted AML12 cells. We used DpnII and obtained over 140 million uniquely aligned read pairs per replicate (Extended Data Fig. 4a). Biological replicates from the same condition were highly correlated Extended Data Fig. 4b) and then we merged replicates for further analysis. We next examined intrachromosomal interactions at different resolutions and found that the Hi-C contact maps in these two samples were similar (Extended Data Fig. 4c). Most of the compartments (97.7%) were unswitched after MATR3 knockdown (Fig. 5a). However, the degree of genome compartmentalization, as visualized by heatmaps of average contacts, showed compartment-specific alteration: increased in AA but decreased in BB compartments (Fig. 5b). Consistently, the interactions between compartment A regions increased but decreased between compartment B regions (Fig. 5c), and the interactions within compartment regions showed opposite trends (Fig. 5d).

**Fig. 5:**
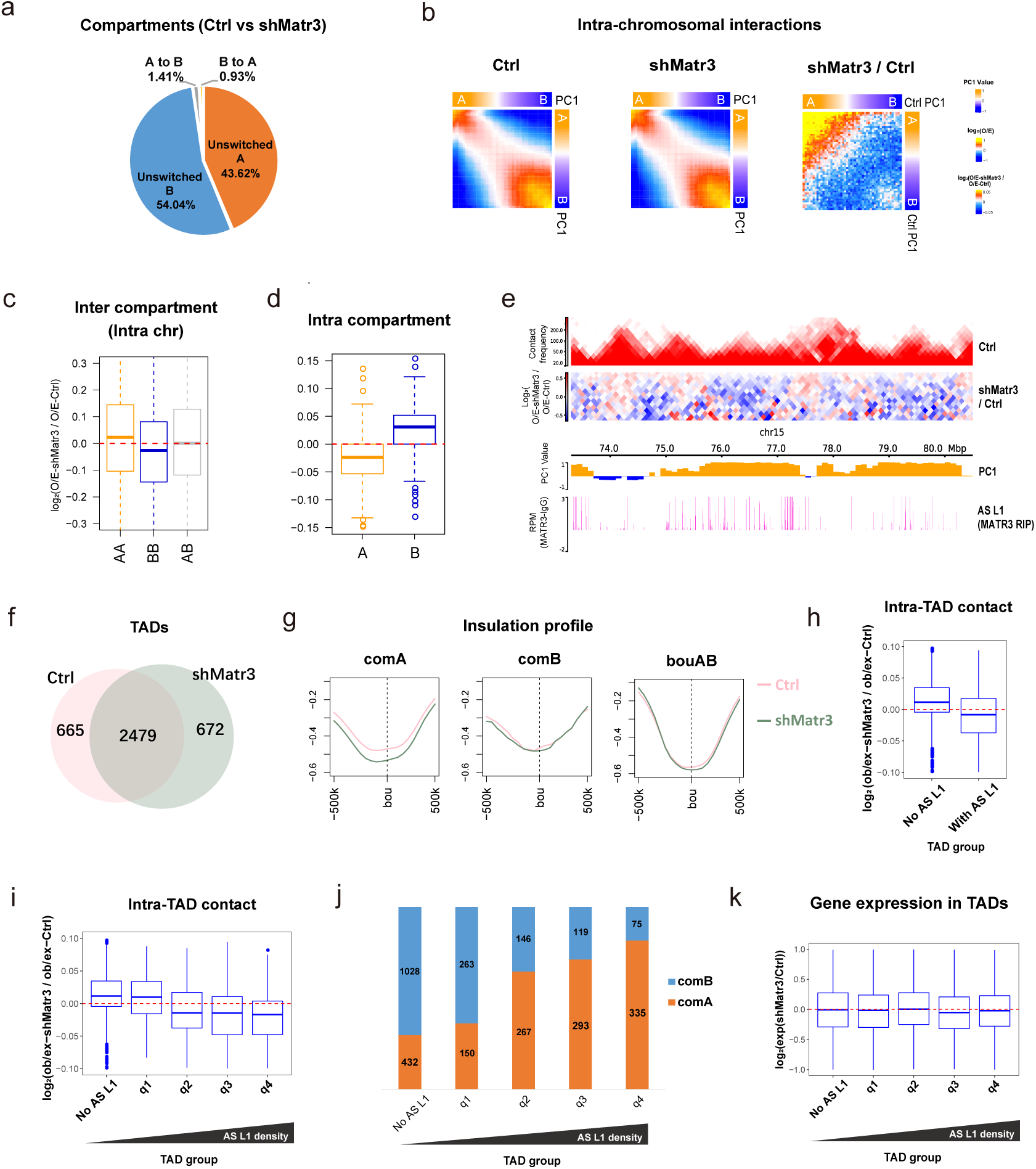
3D genome organization changes on compartments and TADs upon MATR3 depletion. **a**, Percentages of compartment status at unswitched A, unswitched B, B switched to A and A switched to B between Ctrl and shMatr3. **b**, Average contact enrichment between pairs of 100kb loci in Ctrl, shMatr3 and the comparison between them. All the 100kb loci are arranged by Ctrl PC1 values in decreasing order and divided into 50 quantiles. Average enrichment of PC1 values are calculated for each quantile. **c**, Changes in contacts between compartment regions from the same (AA or BB) and different (AB) type in Ctrl and shMatr3. Data are represented as boxplots based on log_2_(O/E-shMatr3 / O/E-Ctrl) values per pair. **d**, Changes in contacts within A or B compartment regions between Ctrl and shMatr3. Data are represented as boxplots based on log_2_(O/E-shMatr3 / O/E-Ctrl) values per compartment region. **e**, Snapshot of an example region, showing Hi-C, AS L1 in MATR3 RIP-seq in Ctrl and Hi-C changes between Ctrl and shMatr3 samples, using HiCExplorer. The values on the y-axis for Hi-C contact and O/E heatmap are iced normalized read counts at 100kb resolution. The values on the y-axis for RIP-seq are average reads per million of mapped reads (RPM). **f**, Venn diagram shows the common and sample specific TADs between Ctrl and shMatr3 samples. The TADs that overlapped length / TAD length > 0.8 both in Ctrl and shMatr3 samples were identified as common TADs. **g**, Insulation strength at boundaries of TAD boundaries in compartment A, B and AB boundaries. **h**, Intra-TAD contacts changes of TADs associated with MATR3-AS L1 RNAs TADs and non-associated TADs according to density of anti-MATR3 RIP-seq signal of AS L1 RNAs. **i**, Intra-TAD contacts changes of TADs non-associated with MATR3-AS L1 RNAs and four quantile groups associated with MATR3-AS L1 RNAs ranking by increasing in MATR3-AS L1 RNAs density. **j**, Percentage of TADs located in compartment A or B regions from five TAD groups (same groups in **i**). **k**, Boxplot shows gene expression changes for TAD groups from **i**.

We also observed the chromatin interaction changes within TADs changed upon MATR3 knockdown (Fig. 5e). We then investigated the Hi-C data at TAD level. After MATR3 depletion, 78.8% TAD boundaries were overlapped between Ctrl and shMATR3 samples (Fig. 5f). By calculating the insulation score, TAD boundaries located in compartment A regions increased in boundary strength, while TAD boundaries located in compartment B regions and AB boundaries hardly changed. (Fig. 5g).

To test whether chromatin interactions at the TAD level were associated with the MATR3-AS L1 meshwork, we compared TADs with or without MATR3-associated AS L1 RNAs, as indicated by MATR3 RIP-seq data. After MATR3 knockdown, the intra-TAD contacts within no-AS L1 TADs increased while decreased within with-AS L1 TADs (Fig. 5h); TADs with the higher AS L1 density exhibited a greater degree of reduction in terms of intra-TAD contacts (Fig. 5i). Furthermore, TADs with the higher AS L1 density were more enriched in compartment A regions, while TADs with no AS L1 density were highly enriched in compartment B regions (Fig. 5j). The overall gene expression changes exhibited no trend along with AS L1 density in TADs (Fig. 5k). Therefore, our Hi-C data suggested that MATR3-AS L1 meshwork confines local clustering of chromatin within TADs in which AS L1 RNAs are highly-transcribed, and most of these regions are in A compartment.

### MATR3-AS L1 meshwork maintain the expression stability of essential genes

We further asked whether the interplay between MATR3 and AS L1 RNAs contributes to gene regulation. We first examined the genomic distribution of the loci that transcribed MATR3-associated AS L1 RNAs. Results showed that 88% of them located in intronic regions of annotated genes (Extended Data Fig. 5a). We then examined genes that contain MATR3-assciated AS L1 on gene body and tested changes of their expression level after MATR3 knockdown. After MATR3 knockdown, the genes containing MATR3-assciated AS L1 were more susceptible to change their expression level (Extended Data Fig. 5b).

Subsequently, to investigate the functional relevance of MATR3-AS L1 meshwork, we examined genes that contain MATR3-assciated AS L1 in intronic regions. MATR3 RIP-seq data in AML12 cells and ES cells were used for these analyses. Genes with AS L1 signals were highly overlapped between the two types of cells (Extended Data Fig. 5c). Gene Oncology (GO) and KEGG enrichment analyses indicated that the common genes were significantly enriched in survival-related pathways including cell cycle, cellular response to DNA damage stimulus and chromatin organization (Extended Data Fig. 5d). This partially explained the observation that cell growth was greatly impeded after MATR3 knock down (Extended Data Fig. 5e). Furthermore, AS L1-associated differentially expressed genes (DEGs) were significantly enriched in terms related to CNS development (neuron projection morphogenesis) and liver function (Cholesterol metabolism with Bloch and Kandutsch−Russell pathways) (Extended Data Fig. 5f).

### Amyotrophic lateral sclerosis-associated MATR3 mutations lead to chromatin redistribution

Multiple mutations of MATR3 were reported to be associated with neurodegenerative diseases including amyotrophic lateral sclerosis (ALS), frontotemporal dementia (FTD), vocal cord and pharyngeal weakness with distal myopathy (VCPDM) and early onset neurodegeneration (EON)^50–52^ (Fig. 6a). Nevertheless, how MATR3 dysfunction contributed to the pathology remained unclear. Aberrant LLPS behavior of neurodegenerative-disease-associated proteins was found to be pathogenic^53^. Therefore, we tested whether mutated MATR3 proteins lead to aberrant phase-separation. Predictor of natural disordered regions (PONDR) algorithm suggested that S85C and F115C, two ALS-associated mutations, could change the PONOR score of MATR3; particularly, the S85C mutation leads a disorder region shifting to an ordered one (Fig. 6b). We thus further investigated the action mechanism of these two MATR3 mutants.

**Fig. 6:**
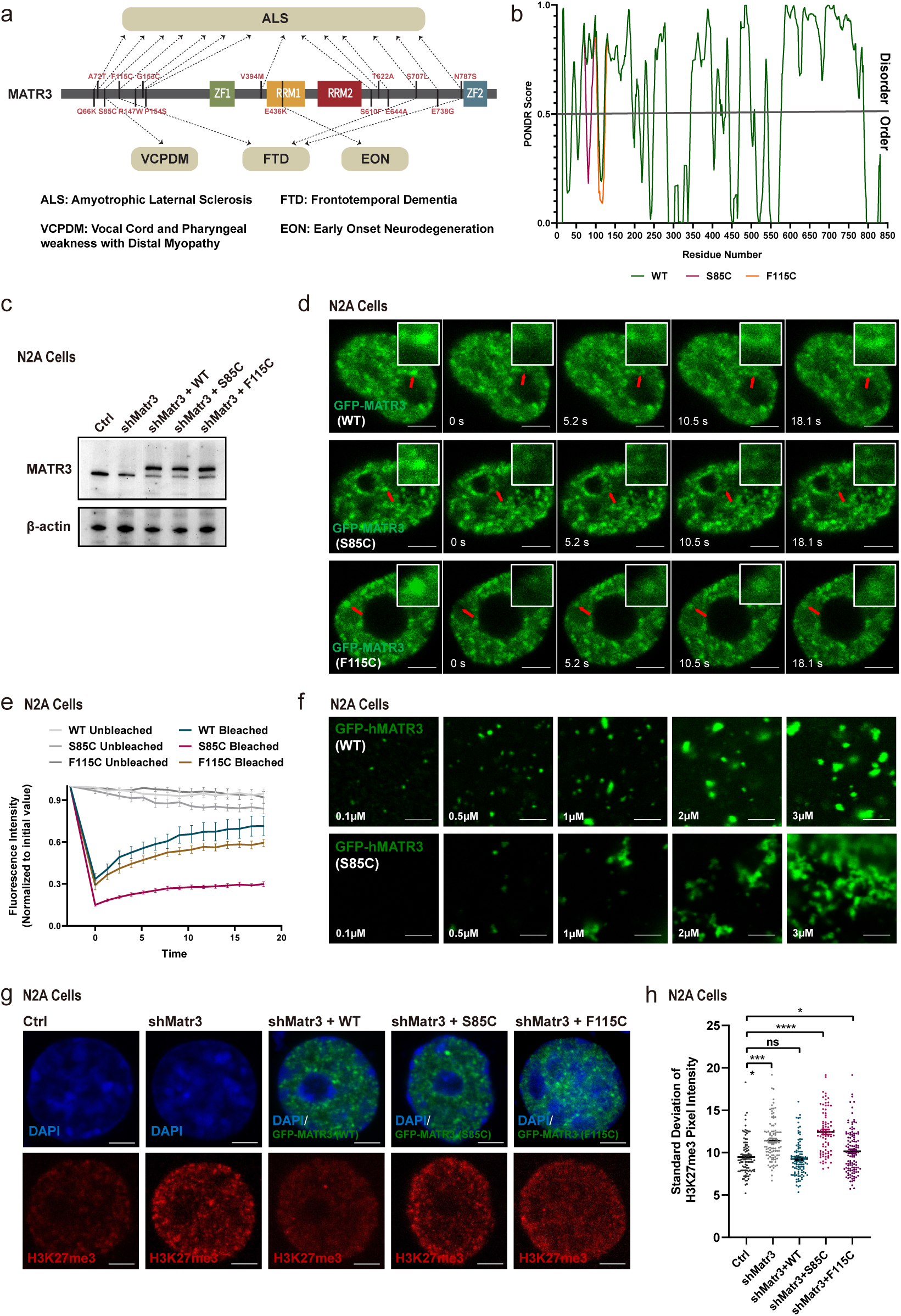
ALS associated mutants lead to reorganization of nuclear chromatin in N2A cells. **a**, Schematic diagram of degenerative-disease-associated mutations on MATR3 protein. **b**, The order/disorder regions of MATR3 (WT/S85C/F115C) protein predicted by the PONDR algorithm. **c**, MATR3 knockdown and GFP-tagged WT/S85C/F115C MATR3 protein replacement in N2A cells. The efficiency of endogenous MATR3 knockdown and exogenous GFP-MATR3 (WT/S85C/F115C) over-expression as detected by western blotting. Representative of two independent replicates with similar results. **d**, Representative images of the GFP-MATR3 (WT/S85C/F115C) FRAP experiments. Solid arrows point to the bleached points. **e**, The fluorescence recovery curve of the GFP-MATR3 (WT/S85C/F115C) FRAP experiments. Data are expressed as the mean ± s.e.m. (n = 9). **f**, Representative images of droplet formation assays with different concentrations of GFP-hMATR3 (WT/S85C) proteins. NaCl concentration, 50mM. Scale bars, 3 μm. **g**, Representative cross-section images showing nuclear localization of H3K27me3 upon Ctrl, MATR3 knockdown and exogenous MATR3 (WT/S85C/F115C) overexpression in N2A cells. Scale bars, 5μm. **h**, Standard deviation of H3K27me3 pixel intensity upon Ctrl (n=93), MATR3 knock down (n=100), GFP-MATR3 (WT) rescue (n=104), GFP-MATR3 (S85C) rescue (n=82) and GFP-MATR3 (F115C) rescue (n=120). The P values were calculated using unpaired two-tailed Student’s t test; ns, not significant, *p<0.05, ****p<0.0001. Error bars indicate mean ± s.e.m. Scale bars, 3μm (**f**) or 5μm (**d**, **g**).

We established a Dox-inducible MATR3 knockdown system in mouse neuroblastoma N2A cells, a common cell model for neurodegenerative disease studies, and then transfected GFP-tagged wild-type or the mutants of MATR3 (S85C and F115C) to replace the endogenous MATR3 protein (Fig. 6c). FRAP assay was used to investigate their dynamic features. We showed that 20 seconds after photobleaching, the fluorescence intensity of GFP-WT, GFP-F115C and GFP-S85C proteins recovered for 57%, 43%, and 20%, respectively, suggesting that these mutations make MATR3 proteins less dynamic in the nucleus (Fig. 6d,e). Furthermore, we purified GFP-tagged human MATR3 WT and S85C proteins and performed *in vitro* droplet formation assay. The WT MATR3 formed into highly dynamic droplets while the S85C MATR3 proteins were much less dynamic and finally assembled into the fiber-like structure (Fig. 6f, **Supplementary Videos. 1,2**). To determine whether the mutation on MATR3 would alter its capacity to interact with AS L1 RNAs, we co-stained MATR3 proteins (WT/S85C/F115C) with AS L1 RNAs in N2A cells. Both wild-type and mutated MATR3 are well-colocalized with AS L1 RNAs in N2A cells (Extended Data Fig. 6a,b).

Finally, we asked whether the MATR3 mutants could change nuclear chromatin distribution. As H3K27me3-modified chromatin showed the highest correlation with MATR3 in N2A cells (Extended Data Fig. 6c,d), we remained to examine H3K27me3 changes in this cell line. After MATR3 depletion, H3K27me3 greatly redistributed in the nucleus with brighter foci appeared; GFP-WT overexpression completely rescued this phenotype. However, neither GFP-S85C nor GFP-F115C rescued the redistribution of H3K27me3 (Fig. 6g,h). These data suggested that ALS-associated MATR3 mutants have changes in their physical state which could further lead to an abnormal chromatin organization.

## Discussion

Based on the data presented, we proposed that MATR3 proteins and AS L1 RNAs phase-separate into a partially dynamic meshwork that facilitate clustering of nearby chromatin (Fig. 7, Left). Mechanistically, upon transcribed, AS L1 RNAs that contain MATR3-binding motifs recruit MATR3 proteins *in-cis*, and the chromatin regions that interact with AS L1 RNAs could be gathered around the meshwork spatially (Fig. 7, Right).

**Fig. 7:**
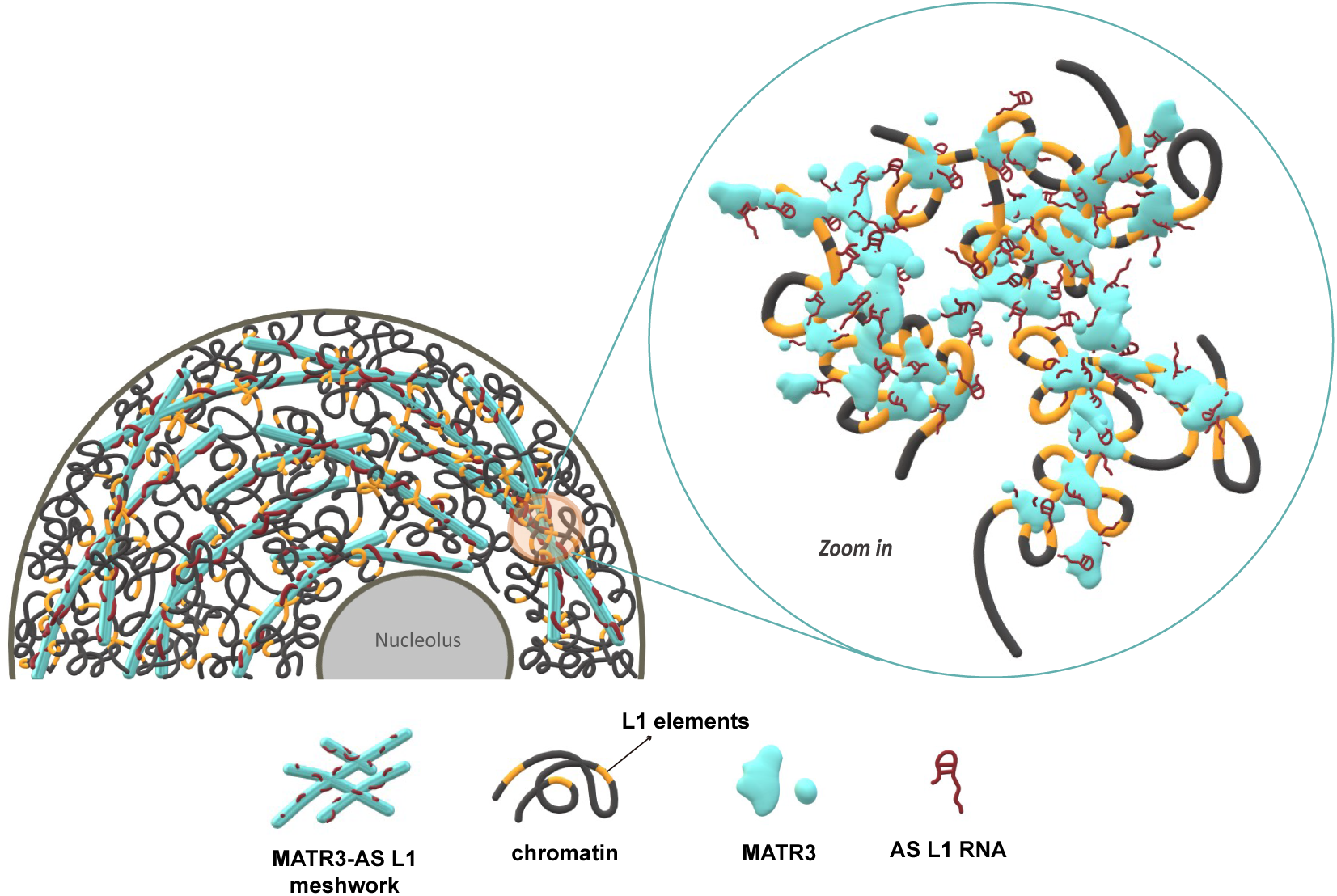
A model on how MATR3-AS L1 RNA meshwork organizes the 3D Structure of chromatin. The meshwork formed by MATR3 proteins and AS L1 RNAs functions as the nuclear scaffold for chromatin which highly transcribed MATR3-associated AS L1 RNAs (Left). The zoom-in view: the newly transcribed AS L1 RNAs *in-cis* interact with L1 loci, attracting MATR3 proteins to form a gel-like meshwork, further gathering nearby chromatin (Right).

NM was observed to be a network in high-salt extracted nucleus since a half century ago^22^. However, there have been few clear microscopic imaging data supporting for existence of NM in living cells, especially the inner NM^54^. Our super-resolution fluorescence microscopy (Fig. 1a) and immuno-electron microscopy (Extended Data Fig. S1a) data showed that MATR3, the representative NM protein, organize as a meshwork and locate on chromatin fibers in the intact nucleus, similar to previous observations in the extracted nucleus^55^. NM is an RNA-protein skeleton and most protein components are RNA-binding proteins^56^. Previous work used to take NM as an entirety, and suggested a consistent mode of action for NM proteins. Our data in this paper and our previous work on SAF-A/HNRNPU and SAFB, performed in the same cell line, indicate that different NM proteins act to regulate different chromatin regions^6, 26^. Their preferential interaction RNAs may account for the discrepancy.

In this paper, we demonstrated the functional roles of AS L1 RNAs in chromatin organization. However, the biogenesis of AS L1 RNAs is still unclear. How do the antisense transcripts be regulated? Are there other partners of AS L1 except for MATR3? What is the function of AS L1 RNAs in development and diseases? Many questions regarding AS L1 RNAs remain to be answered. Previous studies regarding LINEs-derived RNAs seldomly distinguish their transcription orientation^57–59^ or just focused on sense LINEs^15, 17, 18, 20^. The functions and mechanisms of AS L1 RNAs in biological processes have not been well discussed. Based on ENCODE eCLIP datasets, AS L1 RNAs are associated with 9 RBPs (including MATR3, HNRNPM, SUGP2, etc.), and MATR3 eCLIP dataset in HepG2 cells showed the highest AS L1 RNA enrichment^42^. In this study, we revealed that AS L1 RNAs take part in chromatin organization by interplaying with MATR3. It may provide new insights for understanding biological functions of antisense repeat RNAs. Moreover, there are a great number of natural antisense transcripts (NATs) in human cells which is almost equal to sense transcripts, most (>91%) of them are ncRNAs^60^. NATs were suggested to interfere with the expression of sense mRNA through gene silencing, nuclear retention, epigenetic silencing or other mechanisms^61, 62^. Further investigations are needed to reveal the independent role of NATs, especially in the process of 3D genome organization.

Previous work reported that N-terminal of MATR3 could form liquid-like droplets in the cell nuclei of C2C12 mouse myoblasts, suggesting a phase-separation potential for MATR3^63^. This is further proved by the *in-vitro* droplet formation assays in our study. In addition, we demonstrate that MATR3 proteins and AS L1 RNAs comprise a nuclear scaffold, and they function together to maintain a partially dynamic environment for chromatin regulation. There have been a number of instances showing non-coding RNAs and RBPs undergo LLPS which promote cellular sub-compartments formation^64^. In the nucleolus, higher concentration of rRNA may strengthen heterotypic interactions between nucleolus marker proteins (e.g., FBL, NPM1) and increase the nucleolus size^65^. The cytoplasmic lncRNA *NORAD* nucleate droplet formation of Pumilio RBPs (PUM1, PUN2); depletion of *NORAD* leads to dispersal of PUM proteins^3^. These studies revealed that under physiological conditions, higher RNA/protein ratio contributes to larger size of liquid-like phases. However, MATR3 proteins self-organize into liquid-like droplets in cell nuclei when depletion of RNAs; the existence of AS L1 RNAs may abolish the weak interaction between MATR3 proteins and resulted in a gel-like meshwork formation. This indicates that an increase of the AS L1 RNA/MATR3 ratio may facilitate a liquid-to-gel phase transition. We suggest that the gel-like meshwork structure reconciles the dynamics and stability, which is suitable for widespread chromatin regions to switch states in response to spatiotemporal cues.

Recently, knock-in mice models were developed to mimic the two ALS-associated MATR3 mutants (S85C and F115C), and only S85C mice recapitulated pathological features of ALS^52, 66^. In this study, we indicated that S85C shows greater effects on the chromatin mis-localization and protein dynamics than F115C does. A study on the engineered ALS/FTD model mice indicated an abnormal heterochromatin structure and increased staining of H3K27me3 in brain cells^67^. These data suggested that the pathology of S85C mutated ALS may be partially explained by dysregulation of chromatin organization. Our work showed that mutated MATR3 sustains the ability to interact with AS L1 RNAs, but its dynamic activity decreased. Biophysical changes on the MATR3-AS L1 RNA meshwork could decrease their capacity for responding to environmental signals. A previous work in Drosophila model of ALS reported change of L1 RNA expression^68^. Due to the lack of the strand-specific transcriptome datasets for ALS patients (and controls), we could not illustrate the change of MATR3 expression and AS L1 RNA constitution during ALS development. Further investigation focusing on AS L1 RNAs may provide new insights into ALS pathogenesis.

## Methods

### Dox-inducible shRNA system and stable express system in AML12 cells

The mouse hepatocyte cell line alpha mouse liver 12 (AML12; CRL-2254, ATCC, Manassas, VA) were cultured in DMEM/F12 (11320033, Thermo Fisher Scientific, Waltham, MA) supplemented with 10% fetal bovine serum (16140071, Gibco, Grand Island, NY), ITS Liquid Media Supplement (100×, 41400045, Gibco), and 40 ng/ml dexamethasone (D4902, Sigma, Darmstadt, Germany) at 37°C and 5% CO2. For Dox-inducible MATR3-knock down system, plasmids were constructed by cloning the target sequences (The Matr3 shRNA target: GAGACCGATCTTGCTAATTTA) to the pLKO-Tet-On vector ^36^ and then transfect AML12 cells by lentivirus-based system. To generate stable cell lines, AML12 cells were selected in the presence of 1 ug/ml puromycin for 1 day. 1 ug/ml dox was used to induce MATR3 depletion. For stable express cell lines, the GFP-tagged MATR3 cDNA were was cloned into Lv-ef1a-blastsidin-tre-MCS plasmid as described ^69^. GFP-tagged truncations (△ZF1, △ZF2, △RRM1 and △RRM2) were further constructed based on this plasmid using TOYOBO KOD-401 Kit, using primers as described ^70^. Plasmid transfection in AML12 cells was accomplished by lentivirus-based system. To generate stable cell lines, AML12 cells were selected in the presence of 1 ug/ml puromycin and 4 ug/ml blasticidin for 2 days. 1 ug/ml dox was used to induce GFP-MATR3 (or its truncations) expression.

### Rapid protein degradation system in mouse ES cells

The mouse ES cell line (E14TG2a) were cultured in 2i/LIF conditions as described ^71^. The auxin (IAA) inducible MATR3 degradation system was developed in mouse ESCs according to the rapid proteins depletion methods established by Natsume et al ^37^. Firstly, the parental cells were generated by introducing the vector encoding constitutive cytomegalovirus-controlled auxin responsive F-box protein (CMV-OsTIR1) at the safe harbor ROSA26 using CRISPR/Cas ^72^. Cells were further selected in the presence of 1 ug/ml puromycin. After 7 days, colonies were picked for further selection in a 96-well plate and the genotype was checked by genomic PCR. Based on this parental cell line, an in-frame mAID cassette was introduced after the last codon of MATR3 gene by CRISPR/Cas (sgRNA target on Matr3 gene: ATAAATTGGCAGAAGAACGG). Plasmids were transfected with Lipofectamine™ 3000 Transfection Reagent (L3000150, Thermo Fisher Scientific, Waltham, MA). To ensure AID-tagging on both alleles, two short homology donor vectors containing neomycin and hygromycin resistance markers were transfected simultaneously into cells. Cells were further selected in the presence of 2 mg/ml neomycin and 200 ug/ml hygromycin. After 7 days, colonies were picked for further selection in a 96-well plate and the genotype was checked by genomic PCR. 0.5mM IAA (dissolved in alcohol) was used to induce MATR3 degradation.

### Dox-inducible shRNA system and mutant transfection in N2A cells

Mouse Neuro 2a cells (N2A; CCL-131, ATCC, Manassas, VA) were cultured in DMEM (Dulbecco’s Modified Eagle’s medium, Hyclone) supplemented with 10% fetal bovine serum, nonessential amino acids (11140050, Thermo Fisher Scientific) at 37°Cand 5% CO2. Dox-inducible MATR3-Knock down system in N2A cells was established in the same way as in AML12 cells. To generate the shMATR3 resistant MATR3 cDNA, we introduced a synonymous mutation of shMATR3 targeting sites into the Lv-ef1a-GFP-MATR3 plasmid. And based on this plasmid, we further introduced MATR3 mutants (S85C/F115C) using TOYOBO KOD-401 Kit. Lv-ef1a-GFP-MATR3 WT/S85C/F115C (resistant) plasmids were separately transfected into the dox-inducible shMATR3 N2A cells, using standard Polyethylenimine (PEI)-based transfection approach.

### Western Blotting and Immunofluorescence

For western blotting, the cell lysates were blotted against primary antibodies and the blots were visualized with peroxidase-coupled secondary antibodies using ProteinSimple FluorChem M gel imaging system. For immunofluorescence, cells were cultured on the glassy coverslip, fixed by 4% formaldehyde for 10 min, treated by 0.5% Triton X-100 for 10 min, blocked by 4% BSA for 30 min at room temperature, cultured by diluted primary antibody overnight at 4°C, followed by adding second-fluorescence antibody for 1 hour at room temperature and stained with DAPI. Most experiments involving MATR3 antibody were performed by Bethyl (A300-591A), which recognizes carboxy-terminal of MATR3 protein and has higher antibody titers. There was an exception: for detecting the MATR3 degradation efficiency in mouse ES cells. As mAID-tag blocked the antigen-binding sites of the carboxy-terminal MATR3 protein, which could not be recognized by Bethyl (A300-591A), we used the amino-terminal MATR3 antibody (Abcam, ab51081) as an alternative. Other antibody used in this paper as follows: anti-H3K9me3 (ABclonal, A2360), anti-H3K9me2 (ABclonal, A2359), anti-H3K27me3 (ABclonal, A16199), anti-H3K27ac (Active Motif, 39133), anti-H3K4me3 (ABclonal, A2357), anti-βactin (Proteintech, 66009-1-Ig), anti-H3 (Abcam, ab1791).

### RNA fluorescence in situ hybridization (RNA FISH)

RNA fluorescence in situ hybridization was performed using RNAscope® Multiplex Fluorescent Reagent Kit v2 (ACD, 323100). Probes for AS L1 RNA were designed by ACD company (Advanced Cell Diagnostics, Hayward, CA, USA) based on the consensus sequence on antisense L1_Mus1 RNAs. The RNAscope experiment were performed according to the standard RNAscope protocol without protease treating, and then the slides were re-fixed with 4% PFA followed by the standard protocol of immunofluorescence.

### Immuno-electron microscopy

Cells were fixed with 4% formaldehyde and incubated with the primary antibody. The HRP secondary antibody was used to detect the primary antibody and then using DAB (P0203, Beyotime) for the HRP staining, the product is EM-visible. After that, the samples were prepared for conventional transmission electron microscope. In brief, cells were fixed with 2.5% glutaraldehyde, followed by 1% osmium tetroxide treating, gradient dehydration with alcohol, dehydrated with acetone and embedded with resin. The sections were imaged at the EM facility of ION (Institute of Neuroscience, Shanghai, China).

### Chromatin immunoprecipitation sequencing (ChIP-Seq) assay

ChIP experiments were performed as previously described ^73^, antibody against H3K9me3 (Abcam, ab8898), H3K9me2 (Abcam, ab1220), H3K27me3 (ABclonal, A16199) and H3K27ac (Active Motif, 39133) were used. Cells were fixed with 1% formaldehyde for 10 min at room temperature. The libraries were prepared using the VAHTS Universal DNA Library Prep Kit for Illumina V3 (Vazyme, ND607-01) followed by next-generation sequencing (NGS) using the Illumina HiSeq X Ten system.

### Assay for transposase-accessible chromatin with high-throughput sequencing (ATAC-seq)

ATAC-seq experiments were performed as described ^74^. For each sample, 4 × 10^4^ AML12 cells were used. The transposition reaction was incubated at 37 °C for 40 min. The libraries were prepared using the TruePrep DNA Library Prep Kit V2 for Illumina (Vazyme, TD501-01) followed by next-generation sequencing (NGS) using the Illumina HiSeq X Ten system.

### Analysis of subcellular protein fractions

Cells were washed once with 1×PBS and were processed into chromatin-non-associated and chromatin-associated fractions as described ^41^. For RNase A treatment, final concentration of 10μg/ml RNase A (EN0531, Thermo Fisher Scientific) were added to the lysis buffer. The extracts were diluted with equal volume 1×SDS loading buffer and proteins in each fraction were detected by western blotting.

### RNA immunoprecipitation sequencing (RIP-Seq)

RIP experiments were performed as previously described ^75^. Briefly, AML12 cells were cross-linked with UV (4000J), the cell nuclei were extracted and sonicate. Then incubated the supernatant with anti-IgG (CST 2729) or anti-MATR3 (Bethyl, A300-591A) antibodies overnight at 4°C. After wash for 3 times in RIP buffer and elution in 65°C, the RNA samples were extracted with Trizol. Genomic DNA was digested with DNase I (EN0523, Thermo Fisher Scientific) at 37°C for 1h. The libraries were prepared using the VAHTS Total RNA-seq (H/M/R) Library Prep Kit for Illumina (Vazyme, NRM603-01) followed by next-generation sequencing (NGS) using the Illumina HiSeq X Ten system.

### Antisense oligonucleotides (ASOs) treatment

Cells were adherently cultured to about 60% confluency, then the antisense oligonucleotides were added to the culture medium using Lipofectamine™ RNAiMAX transfection regent (13778030, Thermo Fisher Scientific). The final concentration of ASOs is 100mM. ASO target for AS L1 RNA was chose on the consensus sequence on antisense L1_Mus1 RNAs. The following sequences were used: Scramble-ASO (CCUUCCCTGAAGGTTCCUCC)^6^; AS L1-ASO (UAUUGUUGUUUCACCUAUAG).

### Fluorescent recovery after photobleaching (FRAP)

The FRAP was performed as described before ^6^. Briefly, captured one image at pre-bleach, and then an approximately 1mm^2^ region was selected to bleach once with maximum light intensity, following images were captured every 1.3 s on a confocal laser scanning microscopy (Leica TCS SP5, Wetzlar, Germany). Image data analyses was performed using LAS AF Lite software (Leica Microsystems, Wtzlar, Germany).

### *In-vitro* droplet formation assay

**(1)** Protein purification. The GFP-mouse MATR3 (or human MATR3 WT/S85C) cDNA was cloned into the pCAG-flag-6×His plasmid and then transfected into HEK293T cells. The proteins were first purified with Ni Agarose 6 FF (AOGMA) with the AKTA system (GE Healthcare Life Sciences) and further purified with anti-Flag affinity beads. After concentrated in an Amicon Ultracel-50K spin concentrator to exchange the storage buffer [50 mM Tris-HCl (pH 7.5), 200 mM NaCl, 1 mM DTT, 10% glycerol], the proteins were stored in −80°C after flash freezing in liquid nitrogen.
**(2)** *In-vitro* RNA transcription. cDNA of L1, B1 and MajSAT containing T7 promoter sequence and restriction sites hanging on both sides (in opposite orientation) were cloned to the PUC57 vector. And then, linearize the vector with endonuclease for sense- or antisense-oriented transcription, separately. RNAs were transcribed *in-vitro* using TranscriptAid T7 High Yield Transcription Kit (K0441, Thermo Fisher Scientific). UTP in this kit was replaced by Cy5-modified UTP for RNA labeling. The sequences of B1 and MajSAT were obtained from previous reports ^76, 77^ and the representative L1 sequence was a 318bp consensus sequence of L1 Md_F2 elements. **(3)** *In-vitro* droplet formation. Proteins or the mix of protein and RNAs were diluted in PCR tubes at different concentrations in reaction buffer (50 mM Tris-HCl (pH 7.5), 5 mM DTT, 0.1% Triton x-100 and 50 to 800 mM NaCl). Then 3 ul mixed liquid was dropped on the living cell chamber and the images were captured with confocal laser scanning microscopy (Leica TCS SP5, Wetzlar, Germany). Image data analyses was performed using LAS AF Lite software (Leica Microsystems, Wetzlar, Germany).

### In situ Hi-C experiment

The in situ Hi-C libraries were prepared as previously described ^78^. In brief, cells (2.5 millions/sample) were crosslinked by 1% formaldehyde at 25°C for 10 minutes. Then the cell chromatin was digested with 100U MboI at 37°C overnight. Subsequently, filled in the restriction fragment overhangs with biotin at 37°C for 1.5 hours and performed proximity ligation at room temperature for 4 hours. After protein degradation and crosslink reversal, DNA was purified by ethanol precipitation and were sheared to a size of 300-500bp. After size selection, the biotinylated DNA was pulled-down by streptavidin beads and then prepared for Illumine sequencing. The libraries were sequenced via the Illumina HiSeq X Ten system at Annoroad Gene Technology. Two biological replicates were performed for both control and MATR3-depleted AML12 cells.

### Image processing and quantification

The images of immunofluorescence or RNA FISH were obtained with the confocal laser scanning microscopy (Leica TCS SP5, Wetzlar, Germany). To achieve the comparable image data between groups, the microscope parameters were kept unchanged in each set of experiment. To obtain the super-resolution images, we conducted confocal laser scanning microscopy using Leica TCS SP8 STED system, followed by processing the images with a deconvolution software Huygens Ver. Image data analysis was performed using LAS AF Lite software (Leica Microsystems, Wetzlar, Germany) and Image J software (Image J Software, National Institutes of Health, Bethesda, MD, USA).

### ChIP-seq data and ATAC-seq data processing

CutAdapt v.1.16 ^79^ was used to remove adaptor sequences from raw reads of ChIP-seq and ATAC-seq. Reads were mapped to mouse genome (mm10) using Bowtie (v1.2.3) ^80^. Duplicate reads were excluded and kept only one read for each genomic site. BedGraph files were normalized for total mapped read counts using genomecov from bedtools v2.29.2 ^81^. The normalized reads density bigwig tracks were used for visualization with Integrative Genomics Viewer (IGV) ^82, 83^. H3K27me3 peaks were called using SICER ^84^. To generate the ChIP-Seq signal distribution for interested regions, we calculated the average ChIP-Seq signal across these regions.

### RIP-Seq data processing

RIP-Seq reads were mapped to mouse reference genome (mm10) using HISAT2 (v2.1.0) ^85^, with parameters: --rna-strandness RF --dta. Peaks were called using MACS (v1.4.2) ^86^. Read counts were generated using HTSeq-count (v0.6.1) ^87^. BedGraph files were generated using genomecov from bedtools (v2.29.2) ^40^. Integrative Genomics Viewer (IGV) was used for data visualization ^82, 83^.

### Hi-C data analysis

**(1)** Mapping and matrix generation. For Hi-C data, paired-end reads were mapped, processed, and iterative correction (ICE) using HiC-Pro software (version 2.11.4) (https://github.com/nservant/HiC-Pro) ^88^. Read pairs were mapped to the mouse mm10 reference genome (https://hgdownload.soe.ucsc.edu/downloads.html#mouse) with end-to-end algorithm and “-very-sensitive” option. Singleton, multi-mapped, dumped, dangling, self-circle paired-end reads, and PCR duplicates were removed after mapping. To eliminate the possible effects on data analyses of variable sequencing depths, we randomly sampled equal numbers read pairs from each sample for downstream analyses. Valid read pairs were used to generate raw contact matrices at 100-kb, 250-kb and 1-Mb resolutions and applied iterative correction (ICE) on them. We converted .all ValidPairs files to .hic files by the script hicpro2juicebox.sh from HiC-Pro utilities. HiCExplorer (version 3.5.3) (https://hicexplorer.readthedocs.io/en/3.5.3/) ^89^ was used to compute correlations between replicates and plot contact maps for ICE normalized matrices. **(2)** Identification of A and B Compartments. A and B compartments were identified as described previously ^90^. For normalized contact matrix at 100-kb resolution, expected matrix were calculated as the sum of contacts per genomic distance divided by the maximal possible contacts and then converted to a pearson correlation matrix. Then, principal component analysis was performed on the correlation matrix. The first principal eigenvector (PC1) for each bin was used to calculate the overlap ratio with H3K27ac ChIP-seq peaks to assign each bin to A or B compartment. *TAD calling* TADs were called by the insulation score method using cworld software (https://github.com/dekkerlab/cworld-dekker) as previously described ^91^. Insulation scores was calculated by the cworld script “matrix2insulation.pl.” at 100-kb resolution matrix with the parameters “--im iqrMean --is 500000 --ids 250000 --nt 0.3”. The topologically associated domains were identified by the cworld script “insulation2tads.pl” and the 0.3 of min boundary strength was set as the threshold. **(3)** Aggregate contact frequency to compartments. In order to examine the relationship between PC1 values and contacts frequency, we sorted PC1 values for all 100kb bins in decreasing order and divided into 50 equal quantile (Fig. 5b). We then built the 50×50 grid of 2D interval that x axis and y axis are 50 equal quantiles described above. Then in a 50×50 2D interval, we assigned the intra-chromosomal contacts to pairs of 100kb loci in each grid and calculated the average contacts frequency. We then plotted heatmap to show average contacts frequency enrichment in Ctrl and shMatr3 samples and the fold change of contact frequency between them. **(4)** Average contact frequency for compartment regions. To quantify the average enrichment of contacts at compartment level, we first connected the continuous bins that have the same state of PC1 values in Ctrl defined as compartment regions. We then calculated average log ratio of the observed and the expected contacts for A/B intra-compartment regions and inter-compartment regions pairs between the same type (AA and BB) and different types (AB) in Ctrl and shMatr3. Finally, changes in compartment regions between Ctrl and shMatr3 were calculated and represented as boxplots (Fig. 5c,d). **(5)** Insulation score was computed by cworld software (https://github.com/dekkerlab/cworld-dekker)^91^ at 50kb resolution using function matrix2insulation.pl with parameters --is 1000000 --ids 200000 --nt 0.1. **(6)** TADs classified into MATR3-AS L1 RNAs associated TADs and non-associated TADs according to density of anti-MATR3 RIP-seq signal of AS L1 RNAs within TADs (Fig. 5h). MATR3-AS L1 RNAs associated TADs classified into four quantile groups by increasing in MATR3-AS L1 RNAs density (Fig. 5i). GO/KEGG analysis were performed using online software metascape (https://metascape.org/gp/index.html#/main/) (Extended Data Fig. 5d,f)

### Strand-specific RNA-Seq

The libraries were prepared using the VAHTS Stranded mRNA-seq Library Prep Kit for Illumina (Vazyme, NR602-01) followed by next-generation sequencing (NGS) using the Illumina HiSeq X Ten system. Reads were mapped to mouse reference genome (mm10) using HISAT2 v2.1.0 ^85^, with parameters: --rna-strandness RF. FPKM values were calculated using StringTie v1.3.5 ^92^ based on Refgene annotation from the UCSC genome browser, with parameters: --rf. Genes with FPKM >= 1 were considered to be expressed. Read counts were generated using HTSeq-count v0.6.1 ^87^. Differentially expressed genes (DEGs) were calculated using the R package DESeq2 v1.26.0 ^93^, Expression genes were considered differentially expressed if DESeq2 p-value <0.1 and | log_2_(fold changes) | >= BedGraph format files were calculated using genomecov from bedtools v2.29.2 ^81^. Integrative Genomics Viewer (IGV) was used for data visualization in genome ^82, 83^.

### Statistics and reproducibility

Image data analyses was performed using LAS AF Lite software (Leica Microsystems, Wetzlar, Germany) and Image J software (Image J Software, National Institutes of Health, Bethesda, MD, USA). Statistical analyses were performed using GraphPad Prism (8.0) and Microsoft Excel. Representative data with ≥3 independent experiments were expressed as the mean ± s.d. or mean ± s.e.m. Significance testing were accomplished using unpaired two-tailed Student’s t test. No statistical method was used to pre-determine the sample size. No data were excluded from the analyses. The western blots were performed in two or three independent biological replicates with similar results and a representative blot was shown. The ChIP-seq, RIP-seq, ATAC-seq and Hi-C were all performed in two independent biological replicates. The details for each experiment are also provided in the figure legends.

## Acknowledgments

We thank members of the Wen lab for their support and suggestions. We thank Yalin Huang and Jin Li for assistance with fluorescence microscopy. This work was supported by the National Natural Science Foundation of China (32130019 to B.W.) and the National Key Research and Development Program of China (2021YFA1100203 to B.W.).

## Author Contributions

Y.Z. and B.W. conceived and designed this study. Y.Z. performed most of the experiments except for RIP assay (by Z.G) and Hi-C library preparation (Q.W.). X.C. and X.M. performed all bioinformatics analyses, supervised by G.W. X.C, Y.Z. and Z.Z. provided some experimental supports. Y.Z., X.C. and B.W. prepared the manuscript with input of all authors.

---

**Extended Data Fig. 1.**
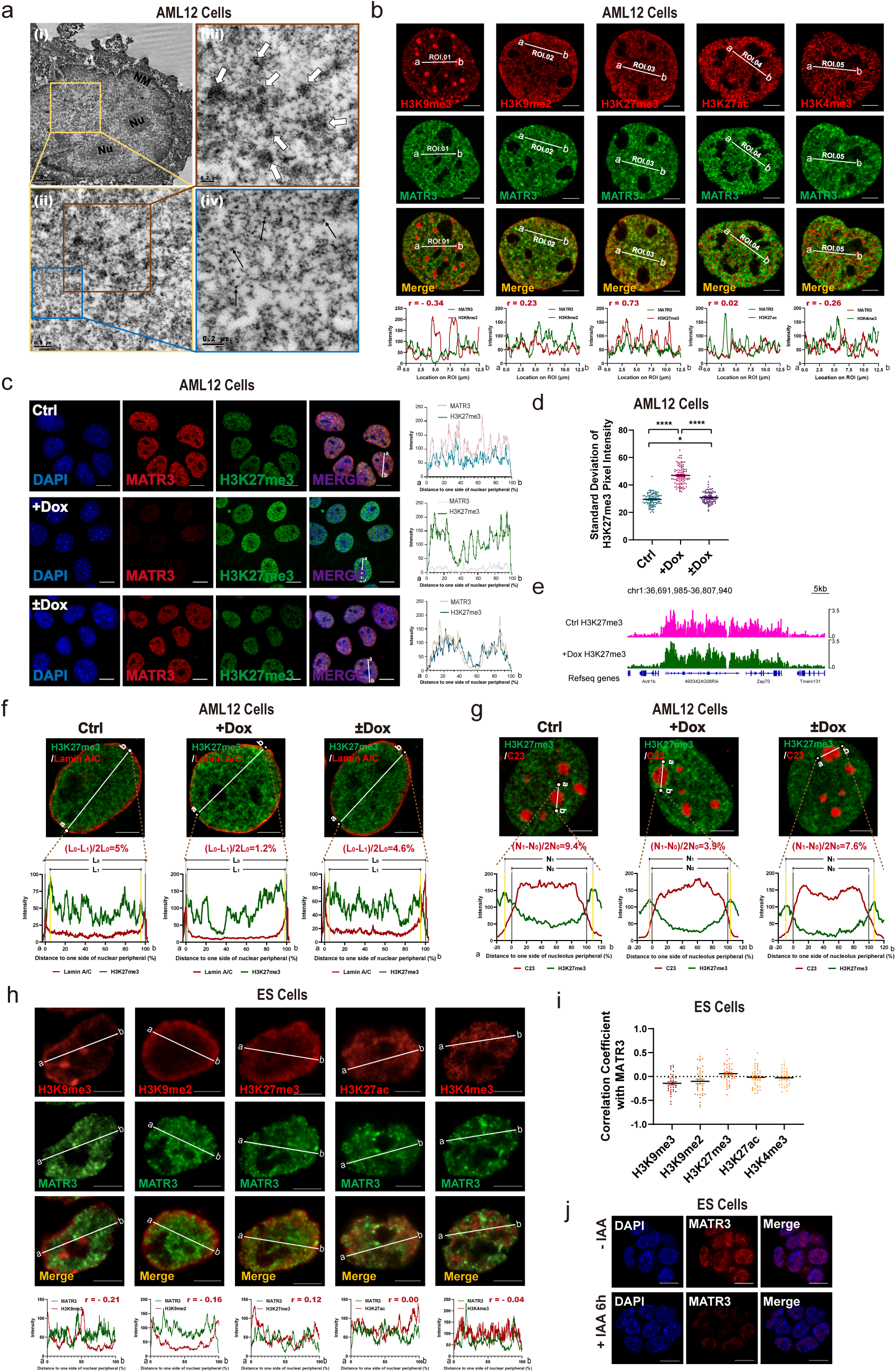
MATR3 modulates redistribution of chromatin in nuclei. **a**, Immuno-electron microscopy analysis of MATR3 protein distribution labeled by DAB in the AML12 cell. The solid arrows point to the individual DAB signal and the hallow arrows point to the clustered DAB signals, both of which represent MATR3 proteins. NM, nuclear membrane; Nu, nucleolus. Scale bars, (i): 2μm; (ii): 0.5μm; (iii), (iv): 0.2μm. **b**, (Upper) Super-resolution fluorescence microscopy images showing relative distribution between MATR3 and histone modifications (H3K9me3, H3K9me2, H3K27me3, H3K27ac and H3K4me3) in AML12 cells. (Lower) Line charts showing pixel intensity of each channel on the regions of interest (ROI). r, coefficient of correlation. **c**, (Left) Representative cross-section images showing nuclear localization of MATR3 and H3K27me3 upon Ctrl, MATR3 knockdown (+Dox) and MATR3 rescue (±Dox). (Right) Line charts showing pixel intensity of each channel on the ROIs. **d**, Standard deviation of H3K27me3 pixel intensity of Ctrl (n=98), MATR3 knockdown (+Dox) (n=100) and MATR3 rescue (±Dox) (n=94). **e**, Genome browser of the H3K27me3 enriched region in Ctrl and MATR3 knockdown (+Dox) samples. **f**, Pixel intensity of ROIs showing relative distribution of H3K27me3 and Lamin A/C upon Ctrl, MATR3 knockdown (+Dox) and MATR3 rescue (±Dox). L_0_, region between nuclear membrane. L_1_, region between two H3K27me3 pixel peaks that closest to the nuclear membrane. Scale bars, 5μm. **g**, Pixel intensity of ROIs showing relative distribution of H3K27me3 and C23. N_0_, region between nucleolus membrane (position of nucleolus membrane on X-axis determined by C23 pixel half-peaks on both sides). N_1_, region between two H3K27me3 pixel peaks that closest to the nucleolus membrane. **h**, (Upper) Representative cross-section images showing relative distribution between MATR3 and histone modifications (H3K9me3, H3K9me2, H3K27me3, H3K27ac and H3K4me3) in ES cells. (Lower) Line charts showing pixel intensity of each channel on the regions of interest (ROI). r, coefficient of correlation. **i**, Coefficient of correlation between MATR3 and histone modification H3K9me3 (n=35), H3K9me2 (n=36), H3K27me3 (n=40), H3K27ac (n=40) and H3K4me3 (n=36) in ES cells. Quantifications were performed on randomly selected ROIs in cell nuclei. Each point represents one cell. **j**, Immunofluorescent detection of the efficiency of MATR3 knockdown in ES cells after 6h addiction of 500μM IAA or equal-volume of alcohol (-IAA). The P values were calculated using unpaired two-tailed Student’s t test; *p<0.05, ****p<0.0001. Error bars indicate mean ± s.e.m. Scale bars, 0.2μm (**a** (iii, iv)), 0.5μm (**a** (ii)), 2μm (**a** (i)), 5μm (**b**, **f**, **g, h**) or 10μm (**c**, **j**).

**Extended Data Fig. 2.**
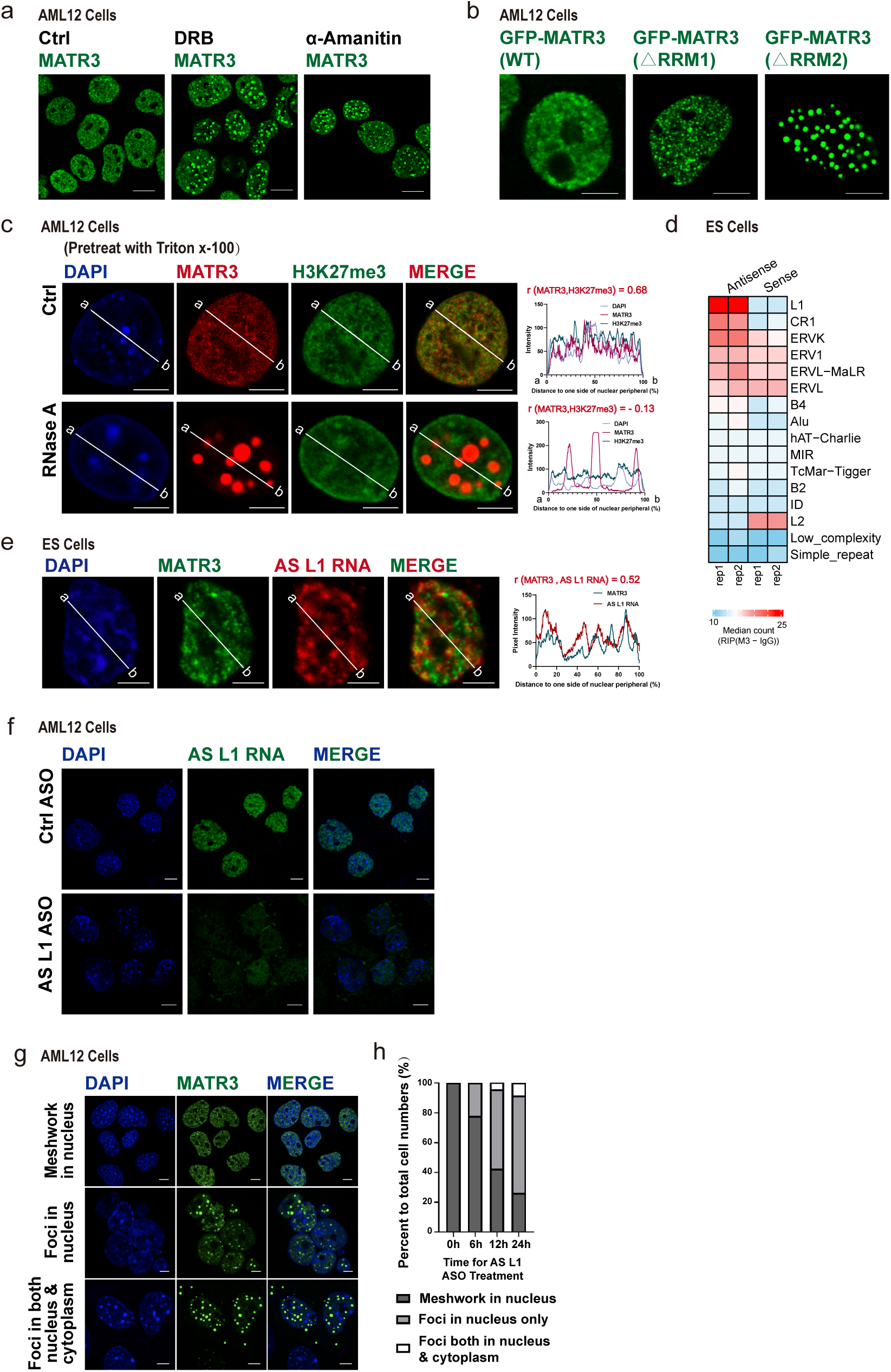
RNAs help maintaining the meshwork structure of MATR3 proteins in nuclei. **a**, The representative cross-section image showing nuclear distribution of MATR3 after 24h treating with 75μM DRB or 24h treating with 50μg/mL α-amanitin in AML12 cells. Scale bars, 10μm. **b**, The representative cross-section image showing nuclear distribution of GFP-tagged MATR3-WT, MATR3-△RRM1 and MATR3-△RRM2. **c**, (Left) The representative cross-section image showing nuclear distribution of DAPI, MATR3 and H3K27me3 before and after RNase A treatment (pretreat with 0.05% Triton x-100 for 30s, followed by 10μg/ml RNase A for 1h) in AML12 cells. Ctrl cells were treated with 0.05% Triton x-100 for 30s. (Right) Line charts showing pixel intensity of each channel on the ROIs. r, coefficient of correlation. **d**, Heatmap of MATR3 RIP-seq sense and antisense median reads count in repetitive elements in ES cells. All RE copies with the RIP (MATR3-IgG) count number >= 10 are kept. Median reads counts are measured for all copies of that RE family. **e**, (Left) Representative cross-section images showing relative distribution between AS L1 RNA with MATR3 in ES cells. (Right) Line charts showing pixel intensity of each channel on the ROIs. **f**, RNA FISH detection of AS L1 RNAs before and after treating with antisense L1 ASOs. **g**, The representative intracellular distribution of MATR3 proteins before and after treating with AS L1 ASOs in AML12 cells. Scales bar, 5μm. **h**, Statistical data for intracellular distribution of MATR3 proteins after 0h (n=1000), 6h (n=1280), 12h (n=1504) and 24h (n=1430) treating with AS L1 ASOs in AML12 cells. Scale bars, 5μm (**a**, **b**, **c**, **e, f, g**).

**Extended Data Fig. 3.**
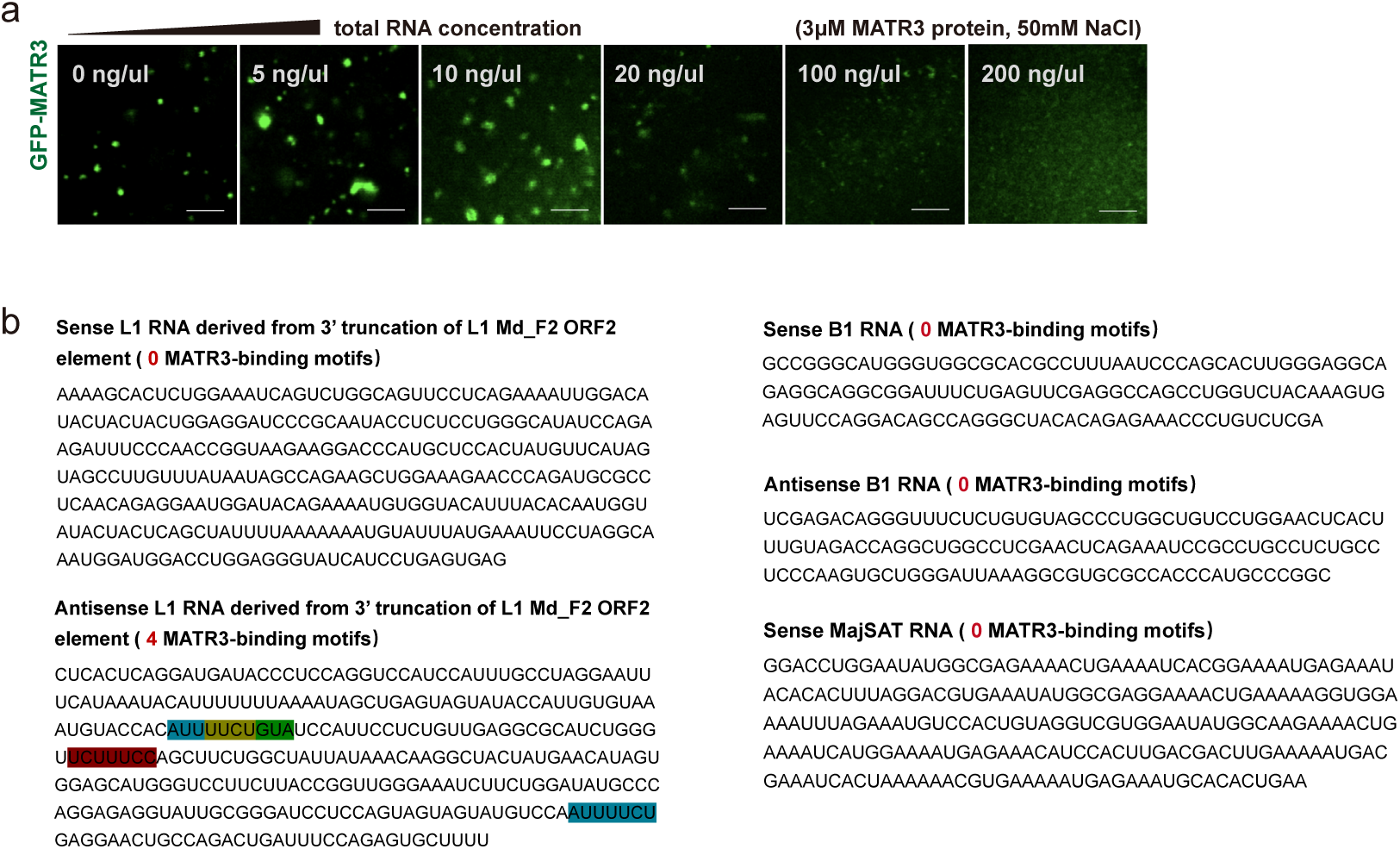
MATR3 proteins interplay with RNAs in vitro. **a**, Representative images of droplet formation assays with 3 μM GFP-MATR3 proteins and different concentration of total RNAs. NaCl concentration, 50mM. **b**, Sequences of sense L1 RNAs, anti-sense L1 RNAs, sense B1 RNAs, anti-sense B1 RNAs and sense MajSAT RNAs used in droplet formation assays. The 7-mer MATR3-binding motifs are highlighted.

**Extended Data Fig. 4.**
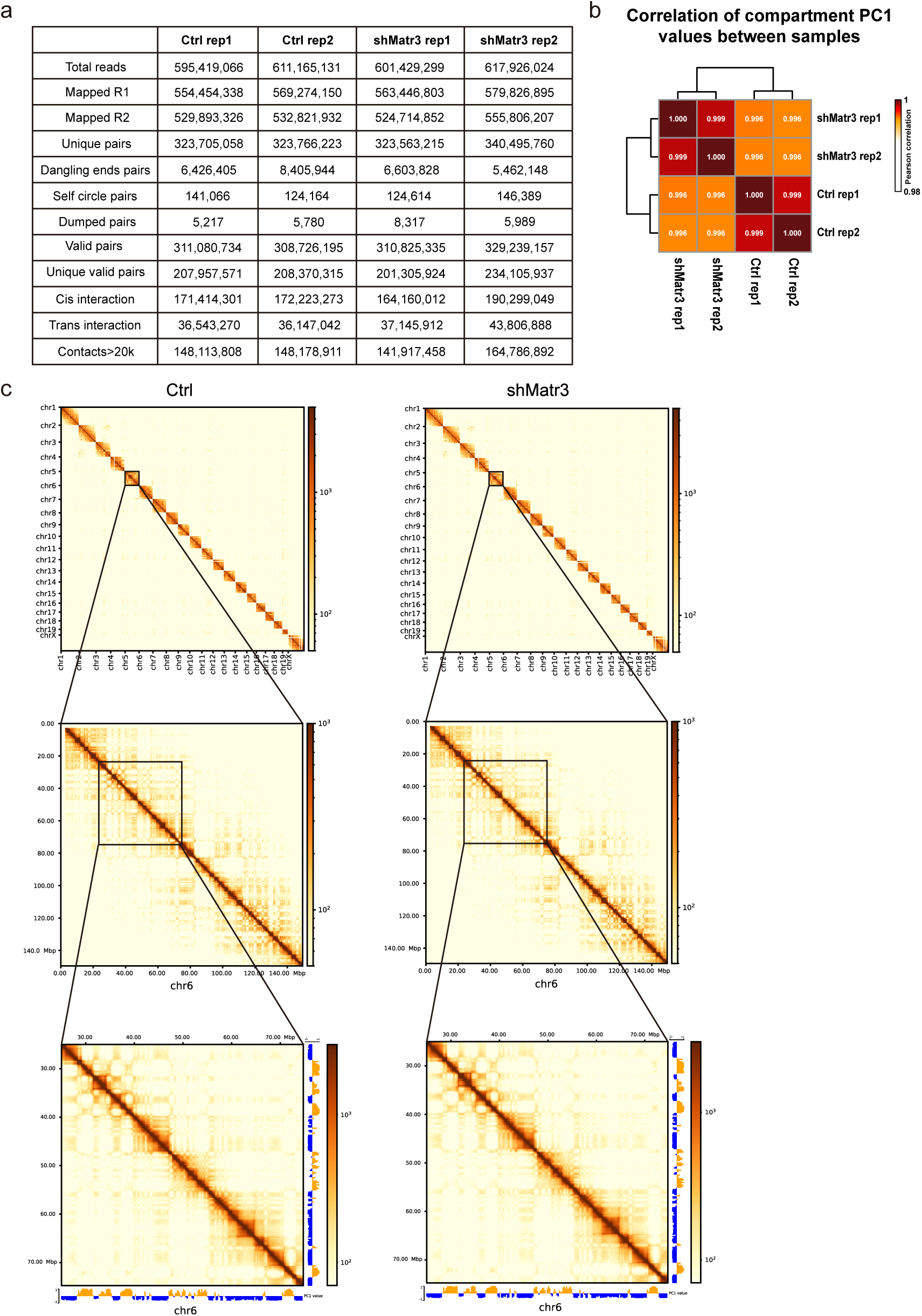
The overall view of Hi-C datasets. **a**, Mapping statistics of Hi-C sequencing data of two replicates in Ctrl and shMatr3. **b**, Pearson correlation coefficients of PC1 values at 250 kb resolution between replicates. **c**, Hi-C contact maps in Ctrl and shMatr3: whole genome at 1MB resolution (top); Chr6 at 250kb resolution (middle); chr6:27-73 Mb at 250kb resolution (down).

**Extended Data Fig. 5.**
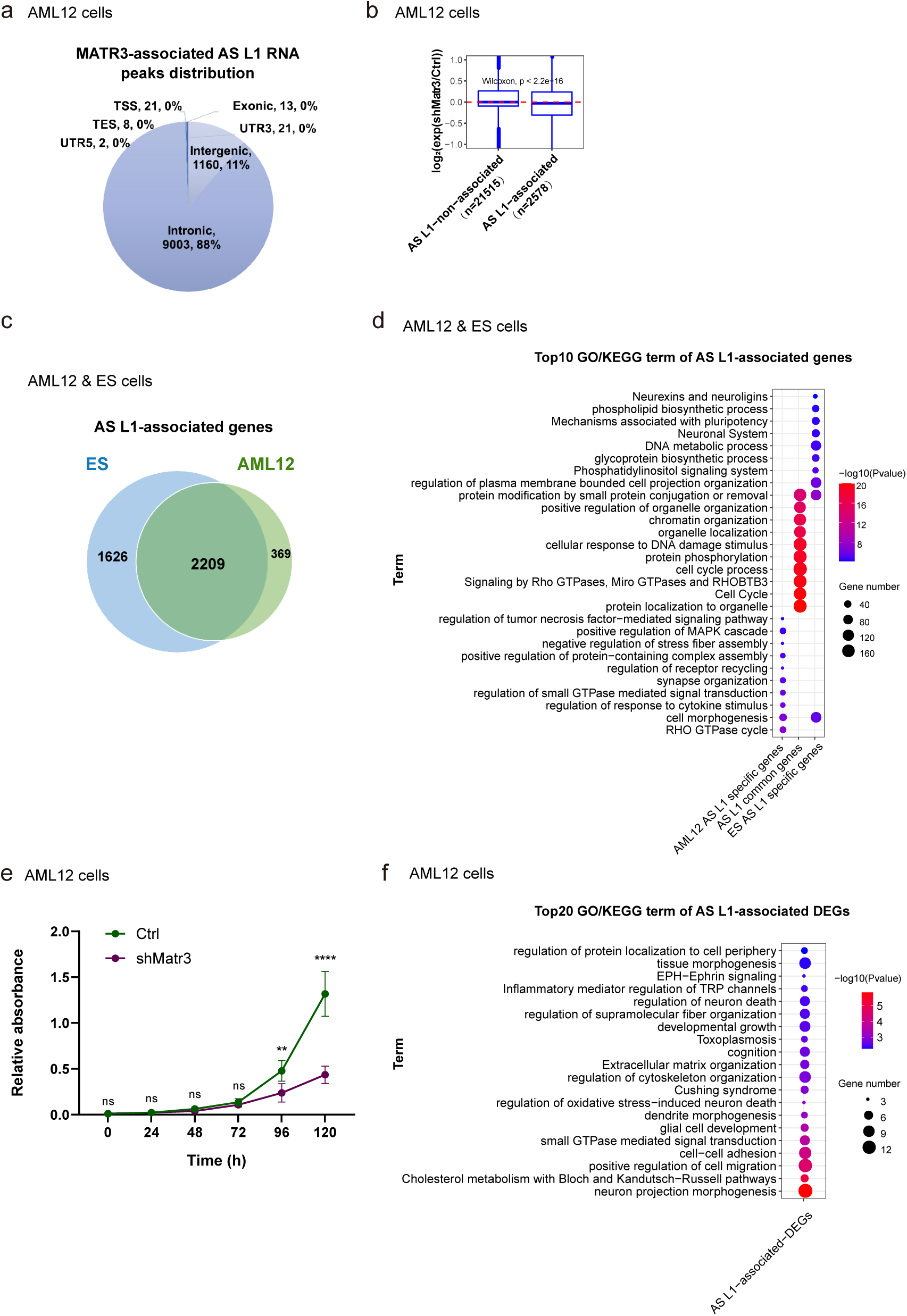
Functional relevance of MATR3-AS L1 RNA associated genes. **a**, Genomic distribution of MATR3-AS L1 RNAs loci in AML12 cells. **b**, Box plot shows gene expression changes in MATR3-AS L1 RNAs associated TADs and non-associated TADs. **c**, Genes that enriched with MATR3-AS L1 RNAs overlap between ESC and AML12 cells. **d**, Top enriched GO/KEGG terms for genes from groups in **c**. **e**, Viability of AML12 cells before and after MATR3 knockdown as detected by CCK-8 assay (n = 7). f, Top enriched GO/KEGG terms for AS L1-associated DEGs (Ctrl vs shMatr3) from AML12 cells. Error bars indicate mean ± s.d.

**Extended Data Fig. 6.**
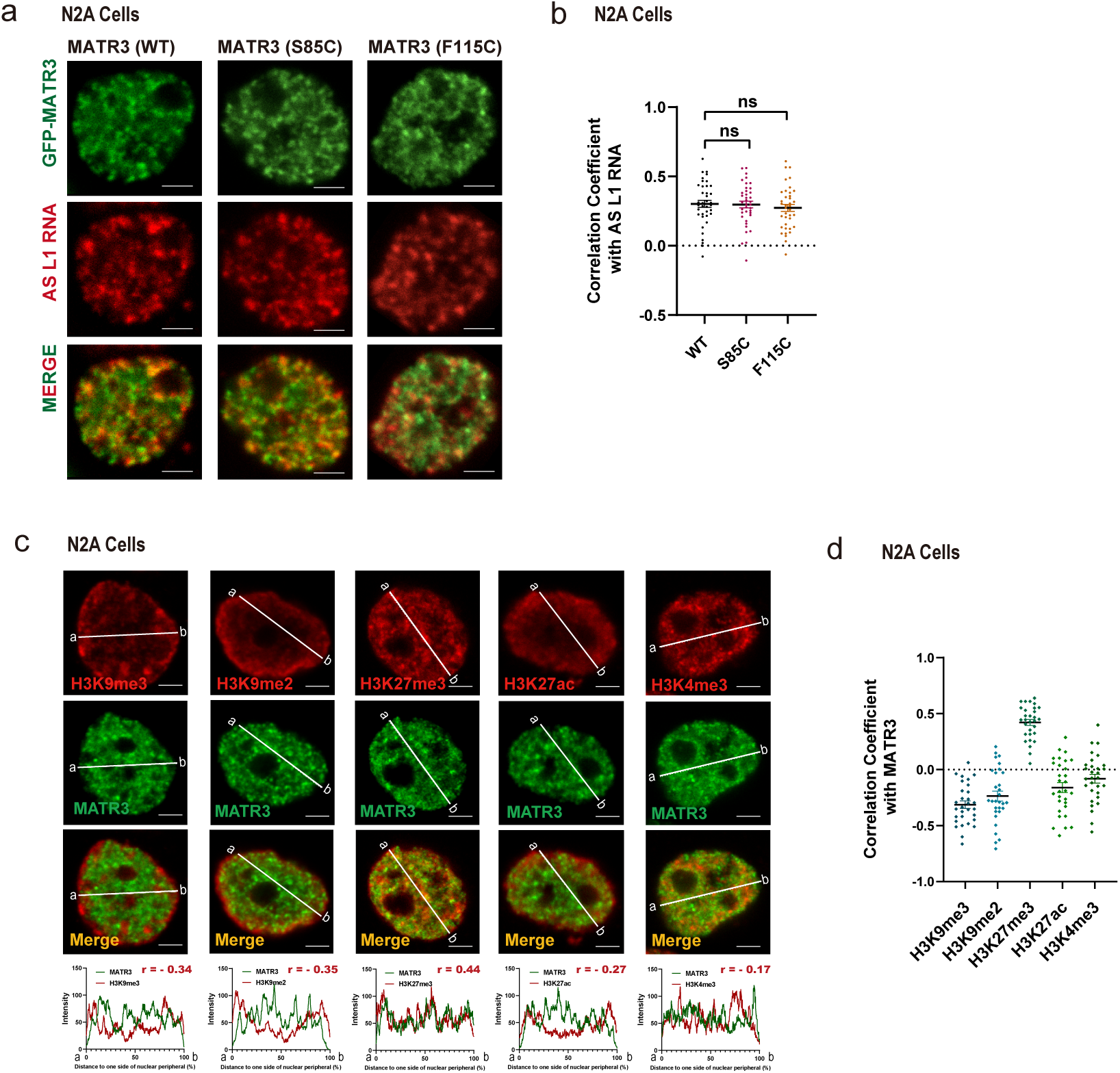
Nuclear distribution pattern of ALS associated mutants in N2A cells. a, Representative images showing nuclear colocalization of AS L1 RNAs with wild-type (WT) and mutant (S85C/F115C) GFP-MATR3 proteins in N2A cells. **b,** Coefficient of correlation between AS L1 RNA with wild-type and mutant GFP-MATR3 proteins. WT (n=38), S85C (n=38), F115C (n=38). The P values were calculated using unpaired two-tailed Student’s t test; ns, not significant. **c,** (Upper) Representative cross-section images showing relative distribution between MATR3 and histone modifications (H3K9me3, H3K9me2, H3K27me3, H3K27ac and H3K4me3) in N2A cells. (Lower) Line charts showing pixel intensity of each channel on the regions of interest (ROI). r, coefficient of correlation. **d,** Coefficient of correlation between MATR3 and histone modification H3K9me3 (n=30), H3K9me2 (n=30), H3K27me3 (n=30), H3K27ac (n=30) and H3K4me3 (n=30) in N2A cells. Quantifications were performed on randomly selected ROIs in cell nuclei. Each point represents one cell. Error bars indicate mean ± s.e.m. Scale bars, 5μm (a, c).

**Supplementary Video. 1. *In vitro* droplet formation video of hMATR3 (WT)**

Representative time-lapse video of droplet formation assay with 3μM GFP-hMATR3 (WT). NaCl concentration, 50mM. Images were captured for 56.760 s at 1.290 s per frame. Scale bar, 2.5 μm.

**Supplementary Video. 2. *In vitro* liquid formation video of hMATR3 (S85C)**

Representative time-lapse video of droplet formation assay with 3μM GFP-hMATR3 (S85C). NaCl concentration, 50mM. Images were captured for 39.990 s at 1.290 s per frame. Scale bar, 2.5 μm.

